# Targeting *TMPRSS2-ERG* Fusion-Driven Prostate Cancer using Novel HBS-α-Helicomimics

**DOI:** 10.64898/2026.10.14.759088

**Authors:** Sushmita Kundu, Promit Ganguly, Lalnunfela Varte, Shifali Mahajan, E.N. Prabhakaran, Bushra Ateeq

**Author notes:** Corresponding Author: Prof. Bushra Ateeq, Molecular Oncology Laboratory, Department of Biological Sciences and Bioengineering, Indian Institute of Technology, Kanpur, Kanpur, 208016, UP, India, The Mehta Family Center for Engineering in Medicine, Indian Institute of Technology Kanpur, Kanpur, 208016, UP, India, Centre of Excellence for Cancer, Gangwal School of Medical Sciences and Technology, Indian Institute of Technology Kanpur, Kanpur, 208016, UP, India (B. Ateeq).

## Abstract

Gene fusions between the androgen receptor (AR)-responsive gene *TMPRSS2* and *ERG* occur in approximately 50% of prostate cancers (PCa) thereby promoting tumorigenesis through aberrant ERG expression. Therapeutic targeting of transcription factor–DNA interfaces remains a formidable challenge, largely owing to the entropic penalty associated with peptide folding during target engagement. Here, we describe the structure-guided design and preclinical evaluation of hydrogen-bond surrogate (HBS)-constrained α-helicomimics targeting the ERG-DNA recognition interface. Structural analysis of ERG-DNA complex revealed that DNA binding induces elongation of recognition helix H4 from approximately 2.5 to 4.0 helical turns. To stabilize the bioactive conformation, we incorporated an N-terminal HBS macrocycle, introduced D363E and S366E mutations to enhance DNA binding and structural stability, and added dual NLS-TAT tags to facilitate cellular delivery. Biochemical and cellular assays demonstrated that the lead compound, ERG helicomimics, competitively inhibited ERG binding to ETS motifs, disrupted ERG association with AR and other cofactors, leading to suppressed ERG transcriptional activity. Chromatin immunoprecipitation-qPCR confirmed reduced ERG occupancy at regulatory regions of canonical target genes, accompanied by downregulation of ERG-driven transcriptional programs. ERG helicomimics also reduced proliferation, migration, and invasion in ERG-positive PCa cells. In a murine xenograft model of *TMPRSS2-ERG*-positive PCa, systemic ERG helicomimics treatment suppressed tumor growth without overt toxicity and reduced Ki-67 expression compared with the designated control peptide. Collectively, our findings support HBS-constrained ERG helicomimics as a rational strategy for targeting ERG-driven PCa and warrant further therapeutic optimization.

## Introduction

Prostate cancer (PCa) is the second most common observed cancer incidence (14.1% of 10.1 million cases) among males worldwide and third (6.1% of 679,421 cases) in India (1,2). PCa is a highly heterogeneous disease associated with several genomic alterations and has been categorized into different molecular subclasses comprising ETS (E26 transformation-specific) family transcription factors)-positive (*ERG, ETV1, ETV4*, *FLI)*, *SPOP*, *CHD1* altered, and *FOXA1* mutated subtype (3-6). Frontline therapy for advanced PCa often involves androgen ablation, which leads to a reduction in ERG expression in patients harboring the ETS gene fusions (7). Androgen independence develops through therapy resistance mechanisms, which restore androgen signaling and *TMPRSS2-ERG* expression. Although a range of drugs targeting androgen biosynthesis or the androgen receptor (AR) have shown efficacy in the treatment of castration resistant PCa, the tumors invariably develop resistance (7).

Notably, transgenic mice conditionally expressing ERG with a *Pten* heterozygous loss background rapidly developed invasive prostatic adenocarcinoma (8,9). Furthermore, knockdown of the *TMPRSS2-ERG* fusion transcript in VCaP cells inhibited cell proliferation, migration, and tumor growth in mice (10-12). Overexpression of ERG transcription factor, has known to induce global changes in chromatin conformation. Genome-wide binding analysis revealed the existence of a highly integrated transcriptional network of ERG and AR (13). Moreover, ERG disrupts AR signaling by depositing repressive histone marks via direct activation of the Polycomb Repressive Complex 2 (PRC2) subunit, EZH2, thereby catalyzing H3K27 trimethylation (13). Another study suggests that the AR and ERG transcriptional complex, together with epigenetic regulators HDACs and EZH2, functions to mediate repression of cytoskeletal genes that promote epithelial differentiation and inhibit metastasis (14). Earlier, our group showed that ∼96% of *TMPRSS2*-*ERG* fusion-positive and ∼70% of AR-positive patients show elevated *DLX1* expression and ERG coordinates with enhancer-bound AR and FOXA1 to drive transcriptional upregulation of *DLX1* in an ERG-positive background (15). Peptidomimetic-based targeting of the ERG transcription factor significantly attenuates proliferation and invasion in *TMPRSS2-ERG* fusion-positive PCa cells, as previously demonstrated (10).

Although *TMPRSS2-ERG*–fusion positive PCa accounts for the predominant molecular subtype, direct pharmacological inhibition of the ERG transcription factor has remained elusive, underscoring a critical therapeutic gap. Current therapeutic strategies largely focus on molecular cofactors and regulators that influence the transcriptional activity of ERG. In this context, our study introduces novel ERG hydrogen-bond surrogate (HBS)-α-helicomimics, which are rationally designed to achieve the desired α-helicity for high-affinity, sequence-selective DNA binding. These helicomimics incorporate cyclic non-peptidic N-terminal HBS-turns, making them thermally stable helical segments, features that enhance their in vivo stability and bioavailability.

This innovative HBS-α-helicomimic technology therefore holds promise as a new class of drug candidates for clinical translation and targeting transcription factors. Given that no direct therapeutic modality has yet been approved for the management of ERG-positive PCa, the development of small peptide inhibitors capable of competitively disrupting ERG-DNA interactions could provide a valuable tool for early-stage intervention in prostate cancer.

## Materials and methods

### Cell lines and culture conditions

Benign prostate epithelial immortalized cell lines (PNT2 (RRID:CVCL_2164) and RWPE1 (RRID:CVCL_3791) and prostate cancer cells (22Rv1 (RRID:CVCL_1045), LNCaP (RRID:CVCL_0395), VCaP (RRID:CVCL_2235), PC-3 (RRID:CVCL_0035), and DU145 (RRID:CVCL_0105)) were obtained from the American Type Culture Collection (ATCC). The cells were cultured and maintained using ATCC-recommended guidelines and media supplemented with 10% fetal bovine serum (FBS) (Gibco) and an antibiotic solution of Penicillin-Streptomycin (Thermo Fisher Scientific), cultured at 37°C and 5% CO_2_. PlasmoTest mycoplasma detection kit (InvivoGen) was used to detect *Mycoplasma* contamination.

### IC50 cell cytotoxicity assay

For determining the IC50 of helicomimics, VCaP cells (3×10^3^) were plated in 96-well dishes and treated with varying concentrations of control peptide and ERG helicomimics (ERG HM-1 and ERG HM-2) for 24 and 48 hours. The IC50 of the helicomimics were determined using WST-1 (Roche). Absorbance was measured at 450 nm using the SpectraMax® Mini Multi-Mode Microplate Reader (Molecular Devices). The data was analyzed using GraphPad Prism 8.43 (RRID: SCR_002798), and IC50 values of the drugs in the cell lines were calculated with the non-linear regression method as previously described (16,17).

### Immunofluorescence

RWPE1-ERG cells were plated in 24-well culture dishes on glass coverslips and treated with control peptide and ERG helicomimics. After 24 h of treatment, cells were fixed with 4% paraformaldehyde in 1X PBS. The cells were washed with 1X PBS, permeabilized using 0.3% Triton X-100 in 1X PBS (PBS-T) for 10 min, and blocked using 5% normal goat serum in PBS-T for 2 h at room temperature. The cells were then incubated with primary antibody (anti-rabbit ERG (1:200, Abcam Cat# ab92513, RRID:AB_2630401) diluted in blocking buffer overnight at 4 C°. Cells were then washed using 0.05% Tween 20 in 1X PBS, followed by incubation with Alexa Fluor conjugated secondary antibodies (1:600 dilution) (anti-mouse or anti-rabbit, CST, Cat# 4408 (RRID:AB_10694704) and Cat# 4412 (RRID:AB_1904025) respectively). The cells were washed again and stained with DAPI (Sigma-Aldrich). The coverslips were mounted on glass slides using Vectashield mounting medium (Vector Laboratories) and sealed on the perimeters. Stained cells were visualized, and images were acquired using a confocal microscope (Nikon, AX/AXR).

### RNA isolation and RT-qPCR

To identify relative target gene expression, total RNA extracted using RNAiso Plus (Takara Bioscience) was converted into cDNA using a First Strand cDNA synthesis kit (Genetix Biotech Asia Pvt. Ltd) according to the manufacturer’s instructions. Quantitative PCR (qPCR) was performed on the QuantStudio5 System (Applied Biosystems) using cDNA template, specific primer sets, and SYBR Green PCR Master-Mix (Genetix Biotech Asia Pvt. Ltd). Primer sequences are provided in Supplementary Table 1.

### Immunoblotting

Cell lysates for immunoblotting were prepared by harvesting the cells in radioimmunoprecipitation assay (RIPA) lysis buffer, supplemented with protease inhibitor cocktail (Genetix) and phosphatase inhibitor cocktail (Calbiochem). The protein samples were prepared in 5X SDS sample loading buffer. The protein samples were separated by SDS-PAGE and later transferred onto a polyvinylidene difluoride (PVDF) membrane (GE Healthcare). Following transfer, the PVDF membrane was blocked by incubating in 5% skimmed milk in 1X TBST (Tris-buffered saline, 0.1% Tween-20) at room temperature for one hour. The blots were then incubated overnight in primary antibody against the protein of interest, diluted in 2% skimmed milk in 1X TBST at 4°C. The primary antibodies include 1:1000 diluted anti-AR (CST, 5153 (RRID: AB_10691711)), anti-ERG (Abcam Cat# ab92513), anti-BRG1 (CST, Cat# 52251, RRID:AB_2799410), anti-FOXA1 (CST, Cat# 58613, RRID:AB_2799548), anti-EZH2 (Abcam Cat# ab191250, RRID:AB_2933983), anti-DLX1 (Thermo Fisher Scientific Cat# PA5-28899, RRID:AB_2546375) and 1:5000 diluted anti-β-Actin (Abcam, ab6276 (RRID: AB_2223210)). The next day, blots were washed with 1X TBST buffer and incubated in horseradish peroxidase (HRP) conjugated secondary antibody (anti-mouse or anti-rabbit antibody, Jackson ImmunoResearch Laboratories, 1:5000 dilution) for two hours at room temperature. Next, the blots were washed with 1X TBST buffer, and the protein signals were visualized using an enhanced chemiluminescence system (ECL, GE Healthcare) according to the manufacturer’s guidelines.

### Cell proliferation assay

Briefly, VCaP and PC3 cells were plated in 96-well culture dishes (3×10^3^ cells/well). At the mentioned time points, cells were treated with control peptide and ERG helicomimics at two concentrations (5µM and 10µM) for 96 h, and the rate of proliferation was determined using cell proliferation reagent WST-1 (Roche). Absorbance was measured at 450 nm using the SpectraMax® Mini Multi-Mode Microplate Reader (Molecular Devices).

### Basement membrane matrix invasion and migration assay

For the cell invasion and migration assay, Transwell Boyden chambers (8μm pore size, Corning) were used. Briefly, the cells were trypsinized, counted, and 1×10^5^ cells were resuspended in 100μl of serum-free media and added to the upper chamber of the Transwell insert coated with Matrigel (Cat. No. 354234, BD Biosciences). In the lower chamber, media supplemented with 20% FBS was added with control peptide and ERG helicomimics. After incubation for 24-48 h at 37°C with 5% CO_2_, the Transwell inserts were gently removed, and a cotton swab was used to remove the Matrigel along with the non-invading cells. The invaded cells on the Transwell membrane were fixed with 4% paraformaldehyde in 1X PBS and subsequently stained with crystal violet solution (0.5% w/v). Representative images were acquired using an Axio Observer Z1 microscope (Zeiss). For quantification, the invaded cells were destained using 10% v/v acetic acid in distilled water, and the absorbance of the destaining solution was measured at 550nm. For the migration assay, a similar protocol was followed, except that uncoated inserts were used (15-18).

### Tumor sphere assay

VCaP cells (2 × 10^4^) were plated in a low-adherence 6-well dish and were cultured in serum-free DMEM-F12 media (1:1, Invitrogen) supplemented with B27 (1X, Invitrogen), EGF (20 ng/ml, Invitrogen), and FGF (20 ng/ml, Invitrogen) as previously described (15). Cells were allowed to grow, and the spheroids formed were collected and mechanically dissociated to form a single-cell suspension before being seeded again in fresh media. The cells were then treated with control peptide and ERG helicomimcs by adding them to the fresh media. Cells were passaged regularly, and peptides were replenished in fresh media at each passage. The experiment was terminated after three weeks, and the prostatospheres were counted and imaged using the brightfield microscope (Leica). The diameter (d) of spheres was quantified using ImageJ (RRID:SCR_003070), and the surface area was calculated using the formula πd^2^. Post-image acquisition, RNA was isolated from these tumor spheres for the quantification of stemness markers.

### Chromatin Immunoprecipitation – quantitative PCR (ChIP-qPCR)

All ChIP experiments were performed as described previously (17). Briefly, crosslinked cells were lysed using lysis buffer (1% SDS, 10 mM EDTA, 50 mM Tris-Cl, and protease inhibitor (Genetix) followed by DNA fragmentation by sonication to an average fragment length of ∼500 bp using an ultrasonicator (Covaris M220 (RRID:SCR_027073)). Sheared chromatin was incubated at 4 °C overnight with 4 µg of primary or isotype control antibodies. Antibodies against ERG (Abcam Cat# ab92513) and control rabbit IgG (Thermo Fisher Scientific Cat# 10500C, RRID:AB_2532981) were used to perform ChIP assay. Simultaneously, the Protein G-coated Dynabeads (Invitrogen) were blocked with BSA (HiMedia) and sheared salmon sperm DNA (Sigma-Aldrich) and incubated at 4 °C overnight. Blocked beads were incubated for 6–8 h at 4 °C with the lysate containing antibody. The antibody-bead conjugates were washed, and the immunocomplex was eluted using elution buffer [1% SDS, 100 mM NaHCO_3_, Proteinase K (Sigma-Aldrich) and RNase A (500 µg/ml each) (Sigma-Aldrich)]. DNA was isolated using the phenol-chloroform-isoamyl alcohol extraction method. Binding sites for the ERG transcription factor on the *DLX1, LDB1*, *CUTL2* and *PLAU* promoters were identified using JASPAR and MatInspector software as previously mentioned (13,15,19). The ChIP-qPCR was performed using primers provided in Supplementary Table 1.

### Luciferase promoter reporter assay

Isogenic VCaP cells were plated in a 24-well culture dish and were transfected with pGL3-DLX1-P wildtype (250 ng) and pRL-null vector (2.5 ng) using FuGENE HD Transfection Reagent. After 48 h of incubation at 37 °C, cells were harvested using the lysis buffer provided with the Dual-Glo Luciferase assay kit (Promega). GloMax® 96 Microplate Luminometer (Promega) was used to measure the Firefly and Renilla luciferase activities according to the manufacturer’s protocol. Renilla luciferase activity was used as a normalization control. The same protocol was followed to measure the luciferase promoter reporter activity for the pGL3-DLX1-P construct (15).

### Whole transcriptome sequencing (WTS)

Two independent replicates of the experiment were performed for each group. After 24 h of treatment with 10μM control peptide and ERG helicomimics for VCaP. Cells were lysed with Trizol and followed by isolation of total RNA using the RNeasy kit (Qiagen). Library preparation and sequencing analysis were performed by Macrogen Inc. (South Korea). Quality of RNA was assessed using a 2100 Bioanalyzer (Agilent Technologies), followed by library preparation (using the Illumina TruSeq Stranded mRNA Library method) and paired-end sequencing on NovaSeq with a 150 bp read length, and more than 25 M reads were recorded for each sample. Samples were subjected to differential expression analysis using DESeq2.

Differential gene expression analysis was performed using the DESeq2 package (1.48.2) (RRID:SCR_000154) (20). Briefly, quality check and preprocessing of WTS reads were performed using FastQC (RRID:SCR_014583) (v0.12.1) and Fastp (RRID:SCR_016962) (v0.23.4)(21). The reads were then aligned to the GRCh38 human reference genome using Hisat2 (v2.2.1) (RRID:SCR_015530) (22) and raw counts were generated using Feature Counts (v2.0.6)(23). Differentially expressed genes were identified using the criteria *abs(log2FoldChange)* > 1 and *padj* < 0.05. Heatmaps were plotted using the ComplexHeatmap R package (v2.25.2) (RRID:SCR_017270) (24), and pathway enrichment was performed and plotted using Metascape (RRID:SCR_016620) (25).

### Mice xenograft study

Athymic nude (Crl:NU(NCr)-*Foxn1^nu^*) (RRID:IMSR_CRL:490), 5–6-week-old male mice procured from Hylasco Biotechnology Pvt. Ltd. (distributor for Charles River Research Models) were anesthetized using a ketamine and xylazine (50 mg/kg and 5 mg/kg, respectively) cocktail injected intraperitoneally. VCaP cells were trypsinized and suspended (2 × 10^6^) in 100 μl of saline with 50% Matrigel and implanted subcutaneously at the dorsal flanks of mice (*n* = 5 for each condition). Tumor burden was measured using a digital Vernier callipers on alternate days, and tumor volume was calculated using the formula (π/6) (*L* × *W*^2^), (*L* = length; *W* = width).

To investigate the anti-tumorigenic effect of ERG helicomimics in a prostate cancer cell-derived xenograft (CDX), 5–6-week-old male athymic nude (Crl:NU(NCr)-*Foxn1^nu^*) mice were subcutaneously implanted with VCaP (2 × 10^6^) cells on both sides of the dorsal flank. Mice were randomized into three groups (*n* = 5 per group) once the average tumor volume reached 75 mm^3^ and treated with control peptide (15mg/kg) and ERG HM-2 (10 mg/kg and 15 mg/kg) administered intraperitoneally for 5 days a week. Tumor burden and mouse weight were measured every alternate day. At the end of the experiment, mice were sacrificed. Tumors were excised for further characterization. All immunodeficient mice used in the study were housed in a specific-pathogen-free (SPF) facility as per the guidelines. The ambient temperature of the mice facility room was maintained at 20–24 °C with ∼40–60% humidity and a 12:12 h light–dark cycle.

### Immunohistochemistry

For Immunohistochemistry (IHC) the tumor tissues excised from the xenografted mice were fixed in 10% buffered formalin, paraffin-embedded, and sectioned at 3-µm thickness using a microtome (Leica). After tissue sections were deparaffinized and rehydrated, sodium citrate buffer was used for heat-induced antigen retrieval (pH 6.0). Furthermore, endogenous peroxidase activity was quenched with 3% hydrogen peroxide (H_2_O_2_) and blocked using 5% normal goat serum. The tissue sections were incubated with the Ki-67 (RRID:AB_2797703) antibody at 4°C overnight. Next morning, the sections were first washed with Tris-buffered saline (TBS) and then incubated with a biotinylated secondary antibody at room temperature for 1 h, followed by incubation with an ABC (avidin–biotin complex, Vector Laboratories) solution for 30 minutes. Sections were then processed for detection of horseradish peroxidase (HRP) activity using 3, 3-diaminobenzidine (DAB) peroxidase (HRP) substrate kit (Vector Laboratories). Quantification for Ki-67 was performed using ImageJ software using 10 random histologic fields.

### Statistical Analysis

For statistical analysis, unpaired two-tailed Student’s t-test, one-way ANOVA, or two-way ANOVA were used unless otherwise specified in the respective figure legend. P ≤ 0.05 was considered significant. The error bars indicate SEM calculated from three independent experiments performed in triplicate.

## Results

### Design and synthesis of ERG helicomimics

Selective competitive disruption of oncogenic ERG transcriptional activity requires an atomic-level understanding of its protein-DNA recognition interface. Structural analysis of the ERG ETS domain (residues 306-419, Chain A) bound to its consensus dsDNA (PDB: 6VGG) revealed a conserved five *α*-helices (H1-H5) and an antiparallel *β*-sheet framework arranged between the N- and C-termini (**Fig. 1A-B**). Helix 4 (H4; residues 362–376) serves as the principal recognition element, inserting into the major groove at the core GGAA ETS motif (**Fig. 1B-C**). The 15-residue in H4 sequence (³⁶²YDKLSRALRYYYDK³⁷⁵N) mediates sequence-specific recognition through an extensive network of hydrogen bonds, salt bridges, and van der Waals interactions (**Fig. 1C-D**). Comparative superposition of free ERG (PDB: 4IRG) with bound ERG revealed a marked induced-fit transition: H4 forms fewer than 2.5 helical turns in the apo state but elongates by approximately 1.0–1.5 turns upon DNA binding, adopting a contiguous ∼4.0-turn helix (**Fig. 1C**). The conformational transition of a flexible linear peptide into the DNA-bound helical state is associated with a substantial entropic penalty; therefore, we employed a HBS scaffold to pre-organize and stabilize the bioactive H4 conformation before DNA binding. (**Fig. 1E-G**).

**Figure 1:**
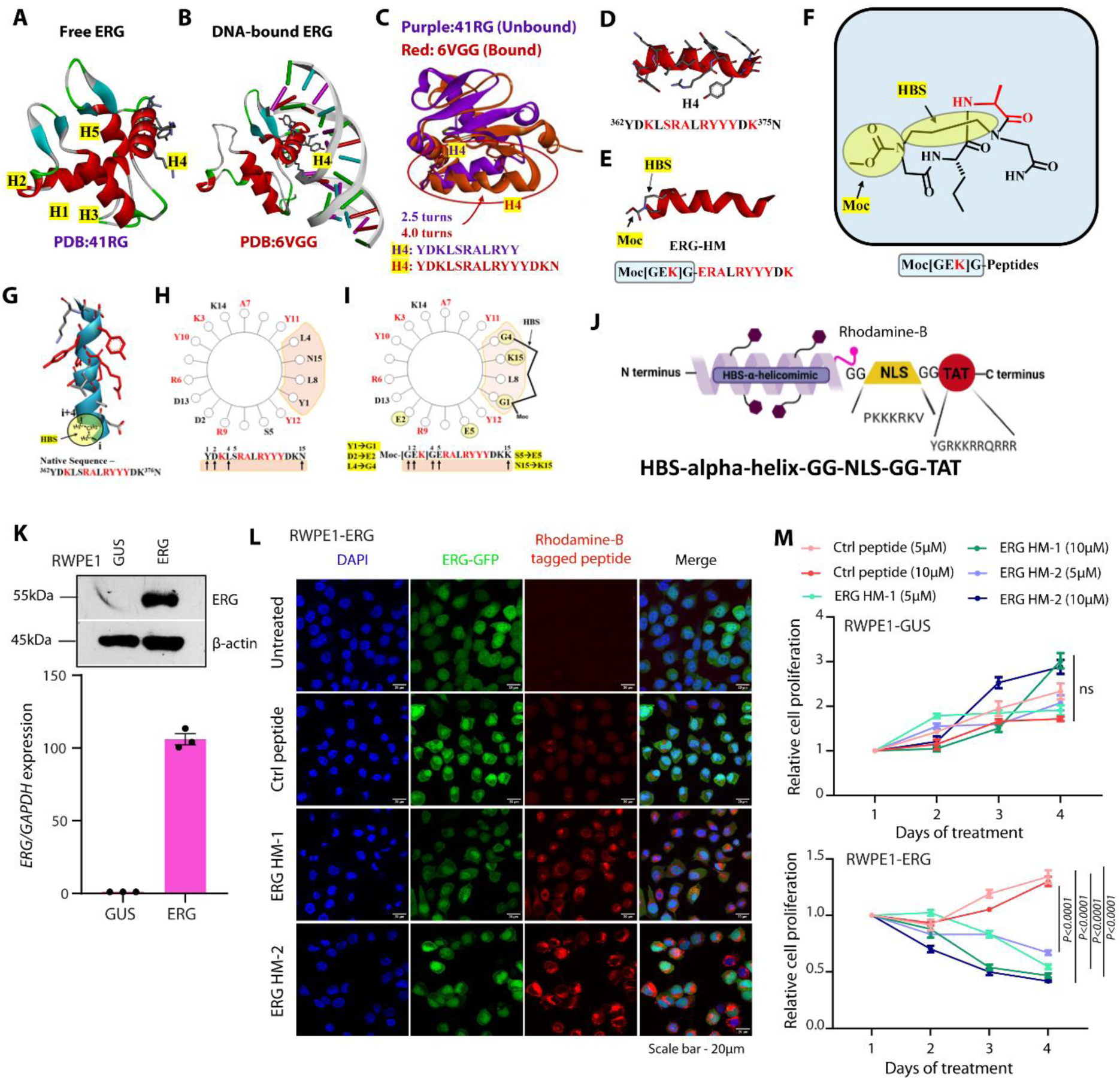
Structure-guided design of HBS-constrained ERG helicomimics and functional evaluation of ERG helicomimics in ERG-expressing prostate cells. **A)** Crystal structure of free ERG (PDB: 4IRG) showing the 2.5-turn recognition helix H4. **B)** ERG DNA-binding domain complexed with target dsDNA (PDB: 6VGG), illustrating major-groove insertion of H4. **C)** Superimposition of unbound (purple, 4IRG) and DNA-bound (red, 6VGG) ERG, highlighting the 1.0–1.5 turn induced-fit elongation of H4 into a 4.0-turn helix. **D)** Native 15-residue H4 with interacting hotspot residues in red. **E-F)** Engineered HBS-constrained ERG helicomimic-(ERG HM-2). **G)** 3D model of the constrained *α*-helical fold. **H-I)** Helical wheel projection of native ERG H4 delineating the non-contact face (shaded yellow) from the DNA-contacting hotspot interface (red). Helical wheel of the engineered analog depicting Gly mutations (Y1G, L4G) accommodating the synthetic HBS crosslink on the non-interacting face, alongside the stabilizing E2---R6 (S366E---R370) salt bridge. **J)** Schematic representation of designed ERG helicomimics. **K)** QPCR and immunoblot of ERG expression in RWPE1-GUS and RWPE1-ERG expressing cell lines. **L)** Immunofluorescence imaging of RWPE1-ERG treated with control peptide and ERG helicomimics. Scale bar: 20µm. **M)** Cell proliferation assay of RWPE1-GUS and RWPE1-ERG expressing cell lines treated with control peptide and ERG helicomimics at 5µM and 10µM.

Distance-based contact mapping of the ERG-DNA complex using Discovery Studio identified the principal residues involved in protein–DNA recognition (**Supplementary Fig. S1**). Helical-wheel analysis showed that the DNA-contacting residues Lys364, Arg367, Ala368, Arg370, Tyr371, Tyr372, Tyr373, and Lys375 cluster on a single amphipathic face oriented toward the major groove, whereas Tyr362, Leu365, Ser366, Leu369, and Asn376 occupy the solvent-exposed, non-interacting face (**Fig. 1H-I**). To stabilize the recognition helix without perturbing the DNA-contacting surface, an HBS macrocycle was introduced across positions 1-4 on the non-interacting face, replacing the canonical i → i+4 backbone hydrogen bond with a propyl crosslink. Tyr362 and Leu365 were replaced with glycine residues to accommodate the macrocycle, generating the N-terminal Moc-[GDK]G-nucleating core in place of YDKL (**Fig. 1I**). In addition, D363E was introduced to extend the side chain toward the major groove and enhance DNA contact, while S366E was incorporated to form an intramolecular salt bridge with Arg370 and reinforce helical stability **(Fig. 1I)** (26-28). For intracellular tracking and nuclear delivery, the optimized helical core was functionalized with a Rhodamine-B moiety at the non-interacting N376K position, followed by nuclear localization signal (NLS; PKKKRKV) and HIV-TAT (YGRKKRRQRRR) sequences. Diglycine spacers were inserted between these modules to provide conformational flexibility and minimize steric interference **(Fig. 1J**). This modular design yielded ERG HM-2: Moc-[GEK]G-ERA-LRYYYDK-K(Rh-B)-GG-PKKKRKV-GG-YGRKKRRQRRR-NH₂ (**Supplementary Fig. S3C**). The second, designated ERG HM-1, was a sequence-reversed analog peptide retaining essential DNA-interacting residues (Ac-KD-YYYRL-ARS-LKDY-K(Rh-B)-GG-PKKKRKV-GG-YGRKKRRQRRR-NH₂) (**Supplementary Fig. S3B**). ERG HM-1 was designed based on published evidence indicating that sequence-reversed analogs can remain biologically active, allowing us to evaluate the contribution of sequence arrangement and pharmacophore orientation to the rational design of peptidomimetic inhibitors (10,29,30).

To distinguish sequence-specific activity from nonspecific effects associated with the delivery sequences, we synthesized one unconstrained analog. A hotspot-disrupted peptide was designated as control peptide-(Ac-YDGL-SGA-LGYGY-DK-K(Rh-B)-GG-PKKKRKV-GG-YGRKKRRQRRR-NH₂) in which key DNA-contacting aromatic and basic residues were substituted to minimize sequence-specific ERG binding while retaining the overall peptide architecture and delivery modules (**Supplementary Fig. S3A**).

ERG HM-2 was synthesized using a convergent solution- and solid-phase strategy. The HBS-constrained helical module was prepared in solution and subsequently coupled to the C-terminal peptide sequence assembled on Rink amide resin. ERG HM-1 and control peptide were synthesized by standard manual solid-phase peptide synthesis (31-35). Following cleavage and global side-chain deprotection, all peptides were purified by reverse-phase HPLC to greater than 98% purity **(Supplementary Fig. S2)**. Scaled purifications were performed using semipreparative or preparative HPLC. Detailed synthetic procedures and full structural characterization (¹H/¹³C NMR and HRMS) are provided in the Supplementary Information

### Cellular uptake and nuclear localization of ERG helicomimics

The designed ERG helicomimics (ERG HM-1 and ERG HM-2) were further modified by attaching a transactivator transcription (TAT) sequence at the C-terminus of the helicomimics to facilitate their internalization into target cells and a nuclear localization sequence (NLS) to ensure their localization into the nucleus of the target cells (10,36), while, Rhodamine B (Rh-B) was tagged onto one of the lysine residues of the helicomimics to enable observation within cells. Additionally, GG spacers were introduced between the helicomimics’ TAT and NLS sequences to prevent functional interference, ensuring that each segment retained its intended role without disrupting the overall activity of the molecule **(Fig. 1J)**. To ensure the internalisation of helicomimics in prostatic cells, we treated the benign immortalized prostate epithelial cell line, RWPE1 overexpressing ERG (RWPE1-ERG) cells and its control RWPE1-GUS, and the ectopic ERG expression was quantified at the transcript and protein levels **(Fig. 1K)**. Immunofluorescence imaging in isogenic RWPE1 showed increased localization of ERG HM-1 and ERG HM-2 inside the nucleus compared to control peptide and as anticipated ERG HM-1 and ERG HM-2 led to diminished ERG fluorescence intensity in RWPE1-ERG cells **(Fig. 1L)**.

### ERG helicomimics inhibit ERG-mediated oncogenicity in prostate cancer

We next determined whether the ERG helicomimics could inhibit ERG-mediated oncogenicity in RWPE1 cells overexpressing ERG. RWPE1-ERG cells treated with ERG HM-1 and ERG HM-2 exhibited significantly reduced proliferation compared to cells treated with the control peptide **(Fig. 1M)**. To extend these findings, we screened a panel of PCa cell lines for ERG expression and identified VCaP as endogenously expressing *TMPRSS2-ERG*-fusion transcript, confirmed at both transcript and protein levels **(Fig. 2A)**. Dose–response cytotoxicity assays demonstrated that the IC50 values of the control peptide were 109.6 µM and 106.0 µM at 24 and 48 hours, respectively **(Fig. 2B-C)**. In contrast, ERG HBS-α-helicomimics displayed much lower IC50 values i.e. ERG HM-1: 10.76 µM (24 h), 11.18 µM (48 h) and ERG HM-2: 16.67 µM (24 h), 17.13 µM (48 h), indicating their cytotoxic effects at lower molar concentrations **(Fig. 2B-C)**. Based on these results, 5µM and 10µM of ERG helicomimics were selected as optimal concentrations for the subsequent functional assays.

**Figure 2:**
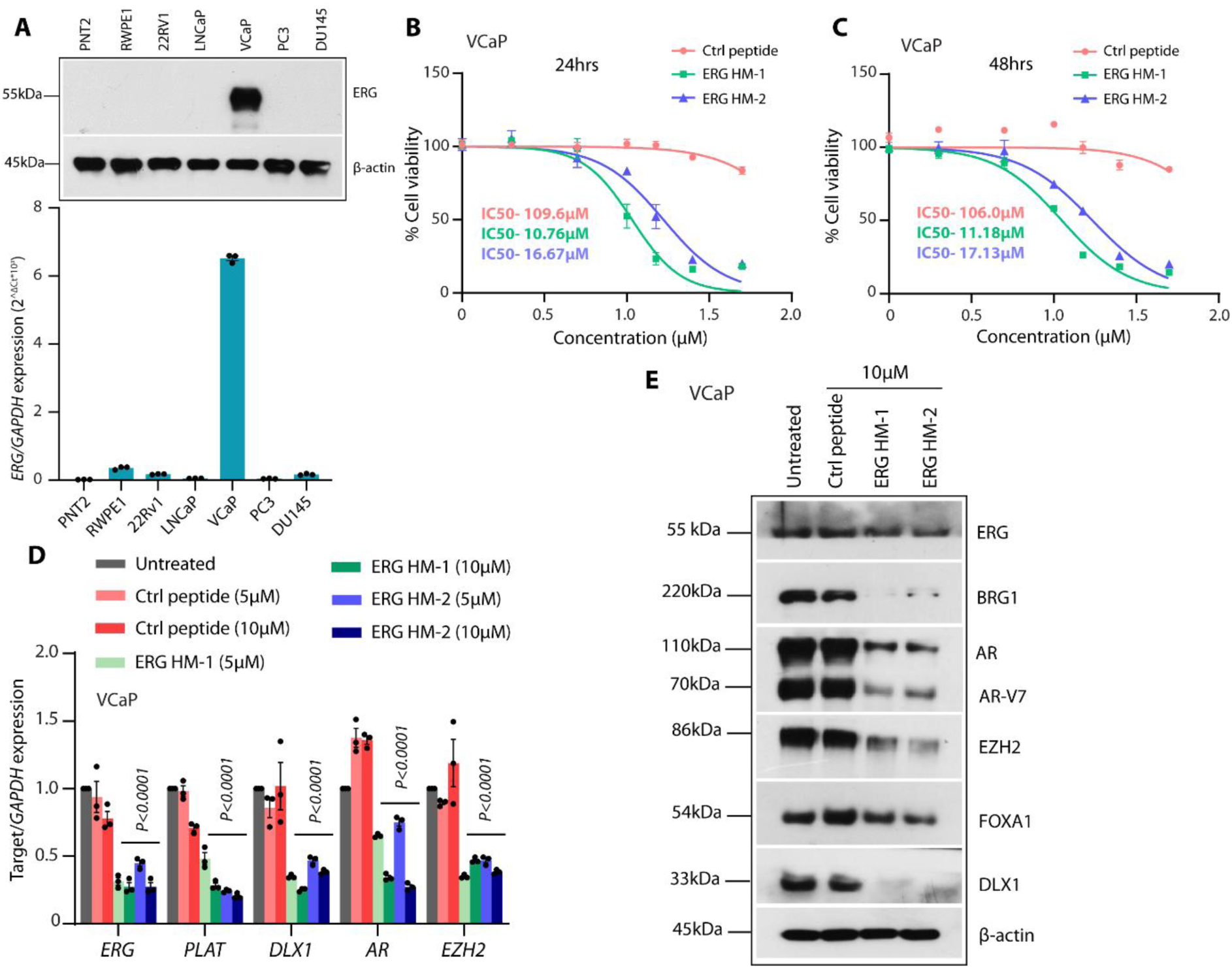
ERG helicomimics reduce VCaP cell viability and downregulate ERG target gene expression in ERG-positive prostate cancer cells. **(A)** QPCR and immunoblot of ERG expression in PCa cell lines. **(B)** Cell cytotoxicity assay to calculate IC50 using VCaP cells treated with control peptide and ERG helicomimics at 24hrs and **(C)** 48hrs with indicated concentrations. **(D)** QPCR of ERG-associated genes in VCaP cells treated with control peptide and ERG helicomimics at 5 µM and 10 µM. **(E)** Immunoblot of ERG target genes in VCaP cells treated with control peptide and ERG helicomimics at 10 µM.

It has been shown that the ERG interacted with AR and DNA-PKcs proteins in a DNA-independent manner (37); thus, we next examined the ability of ERG helicomimics to disrupt these interactions. Quantitative PCR assay demonstrated that ERG helicomimics effectively suppressed ERG-driven transcriptional programs, and decreased transcript levels of canonical ERG target genes, including *PLAT*, *DLX1*, *AR*, and *EZH2*, as well as *ERG* itself **(Fig. 2D)**. Importantly, reduced protein levels of ERG-associated oncogenic regulators AR, DLX1, EZH2, BRG1, and FOXA1 was observed in VCaP cells treated with ERG HM-1 and ERG HM-2 **(Fig. 2E)**. These results highlight the inhibitory potential of ERG helicomimics to attenuate both ERG expression and its downstream oncogenic network.

To examine whether ERG helicomimics would inhibit the ERG-mediated oncogenicity, we performed an array of cell-based assays using VCaP and *TMPRSS2-ERG*-fusion negative PCa cell line, PC3 **(Fig. 3A)**. ERG HM-1 and ERG HM-2 consistently reduced cell proliferation, migration, and invasion in ERG-positive VCaP cells compared with cells treated with the control peptide. In contrast, no significant effects were observed in ERG-negative PC3 cells, indicating the specificity of antitumor activity of ERG HM-1 and ERG HM-2 **(Fig. 3B-C)**. Anchorage-independent growth was further examined by performing the prostatosphere assay, wherein ERG helicomimics markedly impaired both prostatosphere formation efficiency and the prostatosphere area, underscoring their ability to suppress malignant properties **(Fig. 3D)**.

**Figure 3:**
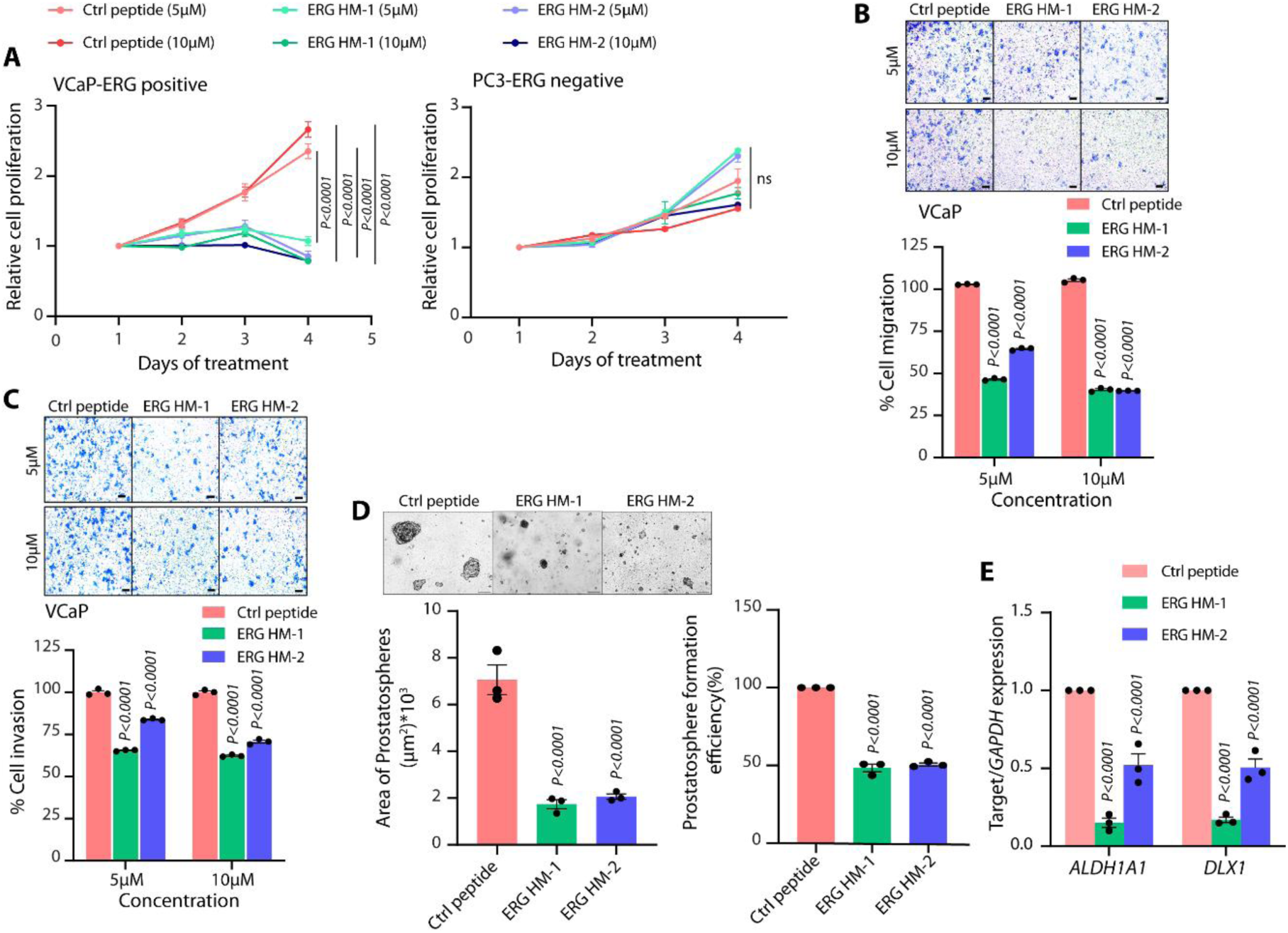
ERG helicomimics selectively inhibit proliferation, migration, invasion, and stemness-associated properties of ERG-positive VCaP prostate cancer cells. **(A-B)** Cell proliferation assay of VCaP (ERG-positive) and PC3 (ERG-negative) cell lines treated with control peptide and ERG helicomimics at 5µM and 10µM. **(C-D)** Boyden Chamber Matrigel cell migration and invasion assay using VCaP cells treated with different concentrations of ERG helicomimics. Representative images for panel **C** and **D** (scale bar 200µm) are shown as insets. (**E)** Bar plot showing area of spheres and number of spheres formed in prostatosphere assay using VCaP under ERG helicomimics treatment conditions. Inset shows a representative image of the spheres formed (scale bar 100µm). Q-PCR data for the *ALDH1A1*, stemness marker and *DLX1* expression using RNA isolated from VCaP prostatospheres. Data represent mean ± SEM. Two-way ANOVA and Tukey’s multiple comparisons were applied.

After termination of the assay, RNA was isolated from the prostatosphere, and the transcript level expression of the stemness marker *ALDH1A1* and the ERG-target gene *DLX1* was also decreased **(Fig. 3E)**. Together, these findings demonstrate that ERG HM-1 and ERG HM-2 disrupt ERG-mediated transcriptional activity, leading to reduced expression of oncogenic protein, and impairing key tumorigenic properties in ERG-positive prostate cancer cells.

### ERG helicomimics attenuates ERG occupancy on its regulatory genes and their transcriptional activity

To determine whether ERG helicomimics could prevent endogenous ERG recruitment to the chromatin, we performed chromatin immunoprecipitation (ChIP) assays using VCaP cells. VCaP cells treated with ERG HM-1 and ERG HM-2 resulted in a marked reduction in ERG occupancy at the promoter of ERG-target genes. ChIP-qPCR analysis revealed a significant decrease in the ERG binding at the *DLX1*, *LDB1*, *CUTL2*, and *PLAU* promoters following ERG HBS-α-helicomimics treatment. In contrast, ERG occupancy remained unchanged in cells that were untreated or treated with the control peptide. *PLAU*, a well-established ERG target gene, served as a positive control and confirmed the specificity of ERG inhibition by ERG HBS-α-helicomimics **(Fig. 4A-D)**.

**Figure 4:**
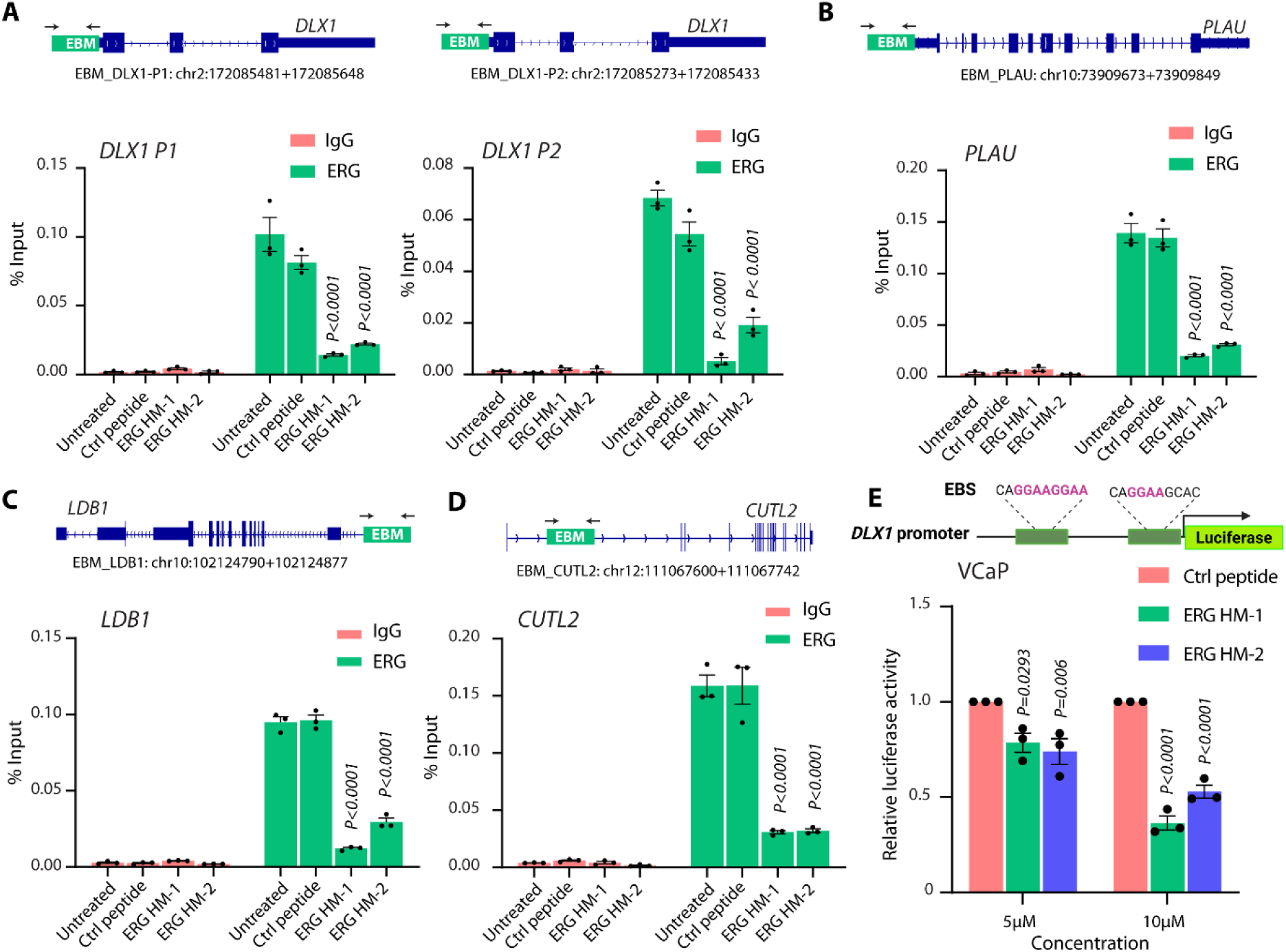
ERG helicomimics disrupt ERG occupancy at target gene promoters and suppress ERG-dependent transcriptional activation of *DLX1* in VCaP cells. **(A-D)** Schema showing chromosomal location of the ERG binding motifs (EBMs) onto *DLX1* promoter selected for ChIP-qPCR (top panel). ChIP-qPCR data showing a decrease in recruitment of ERG at the *DLX1, PLAU, LDB1,* and *CUTL2* promoters respectively in the presence of ERG helicomimics. **(E)** Schema showing the *DLX1* promoter cloned upstream of luciferase gene. Luciferase reporter assay indicating *DLX1* promoter-driven reporter activity in ERG-expressing VCaP cells treated with different concentrations of ERG helicomimics.

To further validate ERG helicomimics-mediated disruption of ERG transcriptional activity, we performed a luciferase promoter reporter assay using the *DLX1* promoter (an ERG target gene) transfected in VCaP cells (15). As speculated, a significant decrease in reporter activity upon treatment with ERG HM-1 and ERG HM-2 was observed in VCaP cells transfected with a *DLX1* promoter reporter, while no significant change was observed with the control peptide **(Fig. 4E)**.

To address ERG helicomimics mediated disruption of ERG recruitment on its target loci, we performed whole transcriptomic sequencing using VCaP cells treated with ERG HM-1, ERG HM-2 as well as control peptide. Several ERG-regulated genes upon treatment with ERG helicomimics were found to be significantly differentially expressed (downregulated) in VCaP **(Fig. 5A-B)**. Genes such as *FASN*, *SREBF1* and *IGF1R* have been implicated in fatty acid biosynthesis, metabolism, cholesterol synthesis **(Fig. 5C-D)** (38-40). Pathway enrichment analysis corroborated our findings as various pathways and gene ontology (GO) terms associated with cholesterol metabolism and homeostasis; SREBF-mediated lipid homeostasis; fatty acid metabolic process; fatty acid and lipoprotein transport were observed to be enriched in the downregulated genesets in VCaP cells treated with ERG HM-1 and ERG HM-2. Further, PCa-risk associated long non-coding RNAs (lncRNAs) such as *PCA3* and *NEAT1* were also found to be significantly downregulated in ERG helicomimics treated VCaP cells (41,42). Additionally, we have identified many protein-coding genes and lncRNAs that have not been investigated for ERG-mediated transcriptional regulation thereby, warranting further research.

**Figure 5:**
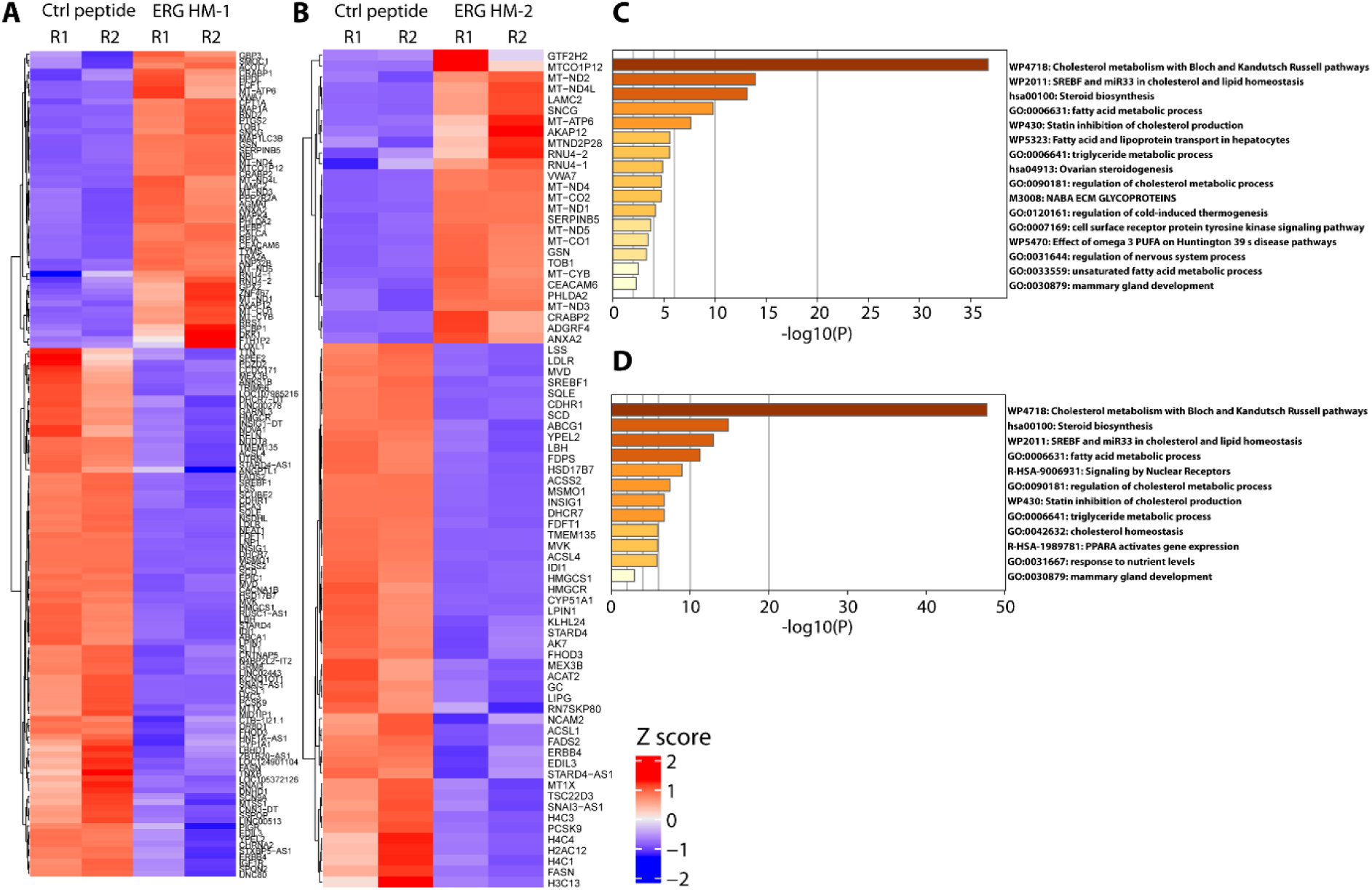
ERG helicomimics alter gene expression and associated signaling pathways in VCaP cells. **(A-B)** Heatmap displaying differentially regulated genes upon ERG helicomimics (ERG HM-1 and ERG HM-2) in VCaP cells compared to control peptide. **(C-D)** Plot showing enriched pathways upon ERG helicomimics treatment.

### ERG helicomimics suppresses ERG-mediated tumor growth in mice

Although ERG HM-1 retained activity against ERG–DNA binding, ERG target-gene expression, and oncogenic phenotype in ERG-positive prostate cancer cells, ERG HM-2 was prioritized as the lead inhibitor in this study since its HBS macrocycle was specifically designed to pre-organize the peptide to achieve a DNA-bound H4 conformation. This conformational restraint preserves the native spatial arrangement of key ERG-DNA contacting residues, reduces the entropic penalty of helix formation, and provides a structural definition for ERG binding motif recognition than the sequence-reversed ERG HM-1. Therefore, to investigate the efficacy of our designed ERG HM-2 against ERG-mediated tumor growth in mouse model, we implanted VCaP cells subcutaneously in athymic immunodeficient male nude (Crl:NU(NCr)-*Foxn1^nu^*) mice. As the tumors reached a palpable stage (average volume ∼75-100 mm^3^), the mice were randomized into three groups (*n* = 5), and the peptides were administered intraperitoneally. Notably, mice treated with ERG HM-2 at two selected doses 10mg/kg and 15mg/kg showed significant tumor regression (∼90%) compared to the mice treated with control peptide **(Fig. 6A-B)**. Subsequently, to delineate the mechanistic impact of ERG HBS-α-helicomimics on tumor xenografts, we performed IHC staining for the proliferative marker Ki-67. Consistent with their proposed mode of action, ERG HM-2 treated groups demonstrated a pronounced reduction in Ki-67 expression **(Fig. 6D)**, reflecting impaired proliferative signaling. These preclinical mouse findings highlight the capacity of ERG-HBS-α-helicomimics to disrupt ERG-driven oncogenic proliferation, thereby reinforcing their therapeutic relevance.

**Figure 6:**
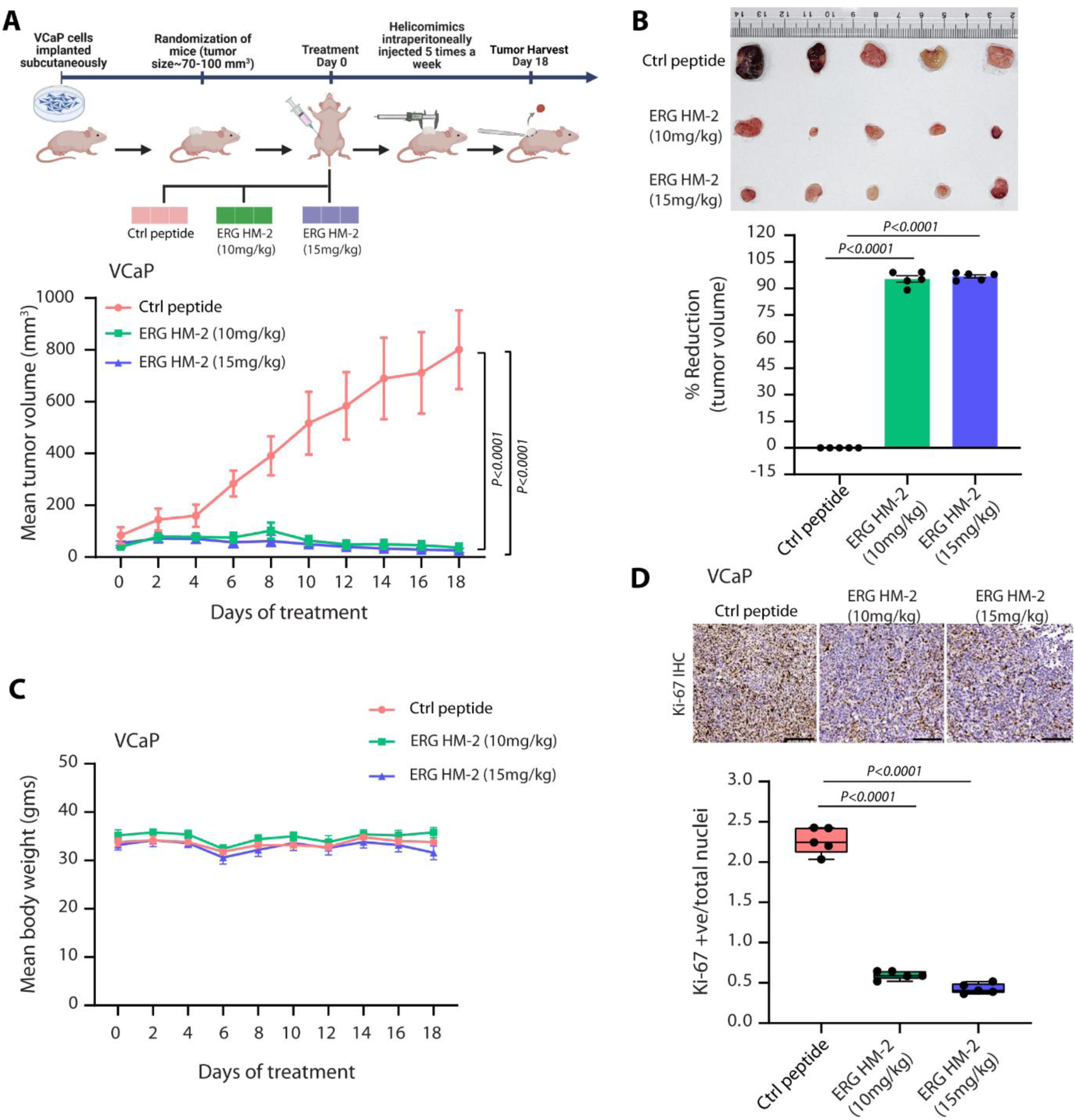
ERG HM-2 inhibit VCaP xenograft growth in vivo. Schematic representation of cell-derived xenograft (CDX) model generation and treatment timeline. ERG-positive prostate cancer cells (VCaP) were injected subcutaneously into athymic nude mice to establish xenografts. Following tumor establishment, mice were randomized into control and treatment groups. ERG HM-2 were administered at selected doses over a three-week treatment period, with tumor volume monitored on alternate days. At the study endpoint, tumors were excised for histological and molecular analyses, including Ki-67 staining to evaluate therapeutic efficacy. **(A)** Mean tumor volume of xenografts generated by implanting VCaP cells in athymic nude mice, and randomized into three treatment groups (*n* = 5 each), namely, control peptide, ERG HM-2 (10 mg/kg), ERG HM-2 (15 mg/kg). **(B)** Bar plot showing percent tumor reduction in the treatment groups (*n* = 5) compared with the control peptide group. **(C)** Mean body weight of mice (*n* = 5 per group) during treatment as mentioned in **A** and **D.** Representative images depicting IHC staining for Ki-67 using formalin-fixed paraffin-embedded tumor xenograft specimens (*n* = 5 per group) as **A**. Scale bar, 50 µm. **(D)** Box plots showing quantification of Ki-67 (*n* = 5) of the mice xenografts as **A**. Data are presented as box-and-whisker plots indicating median (middle line), 25th and 75th percentile (box), and minimum and maximum values (whiskers). For panels **A** and **B**, data represent mean ± SEM. For panels **A, B** and **D**, two-way ANOVA, Tukey’s multiple comparison test.

## Discussion

Resistance to androgen receptor pathway inhibitors (ARPIs) continues to pose a formidable challenge in the treatment of advanced PCa, underscoring the urgent need for therapeutic modalities that circumvent dependence on androgen receptor signalling (7). In this context, targeting ETS transcription factors, particularly ERG, which is aberrantly expressed in approximately 50% of PCa cases owing to *TMPRSS2-ERG* gene fusion, has emerged as a compelling target. However, transcription factors have historically been considered “undruggable” due to the absence of well-defined ligand-binding pockets and their reliance on protein-DNA and protein-protein interactions.

In this study, we developed ERG HBS-α-helicomimic (ERG HM-2) designed to target the ERG oncoprotein by competitively binding to its cognate DNA recognition motifs. This strategy effectively disrupts ERG-DNA interactions, leading to inhibition of ERG-driven transcriptional programs. The selective inhibition of proliferation and invasion in ERG-positive, but not ERG-negative, PCa cells suggests that ERG HM-2 exert their biological effects through specific targeting of ERG-driven signaling pathways. The lack of significant activity in ERG-negative cells supports the on-target mechanism of these compounds and minimizes the likelihood of nonspecific cytotoxic effects. These findings further highlight ERG as a therapeutically relevant vulnerability in ERG-driven PCa and provide evidence that disruption of ERG protein interactions may represent an effective strategy for suppressing tumor progression and metastatic potential. Our findings demonstrate that these ERG-HBS-α-helicomimics attenuate ERG oncogenic activity, thereby establishing their therapeutic potential as a novel modality for targeting ETS transcription factors. Importantly, this approach directly interferes with the transcriptional machinery sustaining tumor progression, offering an advantage over indirect targeting strategies.

These findings are consistent with and extend prior work aimed at inhibiting ERG function. Wang et al. showed that the ERG inhibitory peptides (EIPs) can disrupt ERG–DNA interactions, leading to reduced invasion and tumor growth in preclinical models. The development of retroinverso peptide variants further improved stability and promoted ERG degradation, reinforcing the feasibility of peptide-based ERG targeting (10). Small-molecule inhibitors such as YK-4-279/TK216, VPC-18005, ERGi-USU, and dithiophene diamidines have been explored as alternative approaches, offering potential advantages in pharmacokinetics and systemic delivery (43-46). Recent work by Wang et al. demonstrated that the ERG PNT domain contains a ligandable pocket and identified PBITE-1 as a small molecule capable of directly targeting ERG, providing additional evidence that this undruggable transcription factor can be pharmacologically inhibited (47). While small-molecule targeting of the ERG PNT and ETS domain establishes ERG as a ligandable oncogenic driver, our ERG helicomimic strategy may offer a direct mode of disrupting the protein-DNA interactions underlying ERG transcriptional activity and oncogenicity. Furthermore, mimicking the native helical interaction motif, has the potential to achieve greater specificity and broader inhibition of ERG-dependent transcriptional complexes than approaches that rely solely on occupancy of the protein-protein interactions or degradation. The convergence of these peptide-based and small-molecule strategies highlights a broader paradigm shift, demonstrating that transcription factors like ERG can indeed be therapeutically targeted through innovative chemical and structural design.

Despite these advances, several challenges remain for clinical translation. Peptide and peptidomimetic therapeutics often face limitations related to intracellular delivery, metabolic stability, and potential immunogenicity. Furthermore, given the high degree of structural homology within the ETS transcription factor family, achieving specificity for ERG while minimizing off-target effects on related proteins is essential. Addressing these challenges will require continued optimization of delivery platforms, structural refinement to enhance selectivity, and the development of small-molecule inhibitors based on helicomimic design principles.

From a translational perspective, ERG-targeted therapies are particularly attractive within a precision oncology framework. Diagnostic assays to detect *TMPRSS2-ERG* fusions are already well established in clinical settings, enabling biomarker-guided patient stratification. Consequently, ERG-directed therapeutics could be deployed in a targeted manner, either as monotherapy or in combination with existing treatments such as ARPIs and chemotherapies, particularly in patients with advanced stage or metastatic disease.

In summary, the ERG HBS-α-helicomimic platform provides a strong conceptual and translational foundation for targeting ERG-driven transcription in fusion-positive prostate cancer. By reducing the entropic penalty of helix formation and enhancing local helical stability, this approach demonstrates the importance of structure-guided design in developing effective transcription factor inhibitors. The ability of these helicomimics to disrupt ERG-DNA interactions supports the emerging avenue that oncogenic transcription factors can be pharmacologically targeted. Moreover, the underlying design strategy may be adaptable to other transcription factors implicated in cancer pathogenesis. Collectively, these findings represent an important step toward developing a new class of ERG-directed therapies with potential utility against aggressive ETS-fusion positive prostate cancer.

## Acknowledgements

BA gratefully acknowledges financial support from the Council of Scientific and Industrial Research-ASPIRE grant (37WS(0121)/2023-24/EMR-II/ASPIRE), India Alliance DBT/Wellcome Trust Senior Fellowship (IA/S/19/2/504659); Anusandhan National Research Foundation-J.C. Bose Fellowship (ANRF/JBG/2025/000136/HAA), ANRF-PAIR program (ANRF/PAIR/2025/000002/PAI); Ministry of Education – STARS (MoE-STARS/STARS 2/2023-0219); Department of Biotechnology, Ministry of Science and Technology (BT/PR46959/MED/30/2439/2023), Department of Biotechnology, Ministry of Science and Technology (BT/PR66383/BMS/85/1616/2025) and the Tata Innovation Fellowship (HRD-16012/6/2024-AFS-DBT).

## Authors contribution

B.A. defined the problem. E.N.P. conceived and designed the HBS-helicomimic solution and the control constructs, directed the chemical synthesis. L.V. and S.M. synthesized and purified the helicomimics and control peptides in scales for biological evaluation. S.M., L.V., and E.N.P interpreted the data. S.K. and B.A. conceptualized and designed the biological investigations. S.K. performed the in vitro cell-based assays. S.K. and P.G. executed the whole-transcriptome sequencing and bioinformatics analysis. S.K., P.G., and B.A. conducted the in vivo mouse xenograft studies, performed the statistical analyses, and interpreted the data. B.A. directed the overall biological studies. E.N.P., L.V., S.K., P.G. and B.A. wrote the manuscript. All authors reviewed and approved their respective portions in the final version of the manuscript.

## Ethical approval

Ethical approval for the use of animals in this study was obtained from the Institutional Animal Ethics Committee (IAEC) at the Indian Institute of Technology Kanpur (Approval Number: IITK/IAEC/2023/1201), issued on November 04, 2023, for three years.

## References

1. Sung H, Ferlay J, Siegel RL, Laversanne M, Soerjomataram I, Jemal A, et al. Global Cancer Statistics 2020: GLOBOCAN Estimates of Incidence and Mortality Worldwide for 36 Cancers in 185 Countries. CA Cancer J Clin 2021;71:209–49

2. Mathur P, Sathishkumar K, Chaturvedi M, Das P, Sudarshan KL, Santhappan S, et al. Cancer Statistics, 2020: Report From National Cancer Registry Programme, India. JCO Glob Oncol 2020;6:1063–75

3. Tomlins SA, Rhodes DR, Perner S, Dhanasekaran SM, Mehra R, Sun XW, et al. Recurrent fusion of TMPRSS2 and ETS transcription factor genes in prostate cancer. Science 2005;310:644–8

4. Narod SA, Seth A, Nam R. Fusion in the ETS gene family and prostate cancer. Br J Cancer 2008;99:847–51

5. Kaffenberger SD, Barbieri CE. Molecular subtyping of prostate cancer. Curr Opin Urol 2016;26:213–8

6. Bhatia V, Ateeq B. Molecular Underpinnings Governing Genetic Complexity of ETS-Fusion-Negative Prostate Cancer. Trends Mol Med 2019;25:1024–38

7. Tran C, Ouk S, Clegg NJ, Chen Y, Watson PA, Arora V, et al. Development of a second-generation antiandrogen for treatment of advanced prostate cancer. Science 2009;324:787–90

8. Attard G, Swennenhuis JF, Olmos D, Reid AH, Vickers E, A’Hern R, et al. Characterization of ERG, AR and PTEN gene status in circulating tumor cells from patients with castration-resistant prostate cancer. Cancer Res 2009;69:2912–8

9. Carver BS, Tran J, Gopalan A, Chen Z, Shaikh S, Carracedo A, et al. Aberrant ERG expression cooperates with loss of PTEN to promote cancer progression in the prostate. Nat Genet 2009;41:619–24

10. Wang X, Qiao Y, Asangani IA, Ateeq B, Poliakov A, Cieslik M, et al. Development of Peptidomimetic Inhibitors of the ERG Gene Fusion Product in Prostate Cancer. Cancer Cell 2017;31:532–48 e7

11. Wang J, Cai Y, Yu W, Ren C, Spencer DM, Ittmann M. Pleiotropic biological activities of alternatively spliced TMPRSS2/ERG fusion gene transcripts. Cancer Res 2008;68:8516–24

12. Tomlins SA, Laxman B, Varambally S, Cao X, Yu J, Helgeson BE, et al. Role of the TMPRSS2-ERG gene fusion in prostate cancer. Neoplasia 2008;10:177–88

13. Yu J, Yu J, Mani RS, Cao Q, Brenner CJ, Cao X, et al. An integrated network of androgen receptor, polycomb, and TMPRSS2-ERG gene fusions in prostate cancer progression. Cancer Cell 2010;17:443–54

14. Chng KR, Chang CW, Tan SK, Yang C, Hong SZ, Sng NY, et al. A transcriptional repressor co-regulatory network governing androgen response in prostate cancers. EMBO J 2012;31:2810–23

15. Goel S, Bhatia V, Kundu S, Biswas T, Carskadon S, Gupta N, et al. Transcriptional network involving ERG and AR orchestrates Distal-less homeobox-1 mediated prostate cancer progression. Nat Commun 2021;12:5325

16. Yadav A, Biswas T, Praveen A, Ganguly P, Bhattacharyya A, Verma A, et al. Targeting MALAT1 Augments Sensitivity to PARP Inhibition by Impairing Homologous Recombination in Prostate Cancer. Cancer Res Commun 2023;3:2044–61

17. Khan UK, Goel A, Nigam S, Chaudhary N, Praveen A, Roy A, et al. DKC1 promotes colorectal cancer progression and therapy resistance by dysregulating sphingolipid biosynthesis. Nat Commun 2026;17

18. Manzar N, Khan UK, Goel A, Carskadon S, Gupta N, Palanisamy N, et al. An integrative proteomics approach identifies tyrosine kinase KIT as a therapeutic target for SPINK1-positive prostate cancer. iScience 2024;27:108794

19. Bharti M, Bajpai A, Rautela U, Manzar N, Ateeq B, Sinha P. Human ERG oncoprotein represses a Drosophila LIM domain binding protein-coding gene Chip. Proc Natl Acad Sci U S A 2023;120:e2211189119

20. Love MI, Huber W, Anders S. Moderated estimation of fold change and dispersion for RNA-seq data with DESeq2. Genome Biol 2014;15:550

21. Chen S, Zhou Y, Chen Y, Gu J. fastp: an ultra-fast all-in-one FASTQ preprocessor. Bioinformatics 2018;34:i884–i90

22. Kim D, Paggi JM, Park C, Bennett C, Salzberg SL. Graph-based genome alignment and genotyping with HISAT2 and HISAT-genotype. Nat Biotechnol 2019;37:907–15

23. Liao Y, Smyth GK, Shi W. featureCounts: an efficient general purpose program for assigning sequence reads to genomic features. Bioinformatics 2014;30:923–30

24. Gu Z, Eils R, Schlesner M. Complex heatmaps reveal patterns and correlations in multidimensional genomic data. Bioinformatics 2016;32:2847–9

25. Zhou Y, Zhou B, Pache L, Chang M, Khodabakhshi AH, Tanaseichuk O, et al. Metascape provides a biologist-oriented resource for the analysis of systems-level datasets. Nat Commun 2019;10:1523

26. Olson CA, Spek EJ, Shi Z, Vologodskii A, Kallenbach NR. Cooperative helix stabilization by complex Arg-Glu salt bridges. Proteins 2001;44:123–32

27. Shi ZS, Olson CA, Bell A Jr, Kallenbach NR. Stabilization of α-helix structure by polar side-chain interactions: Complex salt bridges, cation-π interactions, and C-H•••O H-bonds. Biopolymers 2001;60:366–80

28. Meuzelaar H, Vreede J, Woutersen S. Influence of Glu/Arg, Asp/Arg, and Glu/Lys Salt Bridges on alpha-Helical Stability and Folding Kinetics. Biophys J 2016;110:2328–41

29. Cerchietti LC, Yang SN, Shaknovich R, Hatzi K, Polo JM, Chadburn A, et al. A peptomimetic inhibitor of BCL6 with potent antilymphoma effects in vitro and in vivo. Blood 2009;113:3397–405

30. Snyder EL, Meade BR, Saenz CC, Dowdy SF. Treatment of terminal peritoneal carcinomatosis by a transducible p53-activating peptide. PLoS Biol 2004;2:E36

31. Kumar A, Prabhakaran EN. Thermal Folding of Peptides into Single Helical Turns. Chembiochem 2025;26

32. Pal S, Banerjee S, Prabhakaran EN. Helix-Coil Transition at a Glycine Following a Nascent alpha-Helix: A Synergetic Guidance Mechanism for Helix Growth. J Phys Chem A 2020;124:7478–90

33. Pal S, Banerjee S, Prabhakaran EN. alpha-Helices propagating from stable nucleators exhibit unconventional thermal folding. FEBS Lett 2021;595:2942–9

34. Prabhakaran EN, Pal S, Kumar A, Mahajan S. Large Scale Synthesis of Native Turn and Helix Mimics Stabilized by a Generic Hydrogen Bond Surrogate. 2021

35. Reddy SS, Pal S, Ghosh S, Prabhakaran EN. Hydrogen Bond Surrogate-Constrained Dynamic Antiparallel beta-Sheets. Chembiochem 2021;22:2111–5

36. Bajpai A, Quazi TA, Tang HW, Manzar N, Singh V, Thakur A, et al. A Drosophila model of oral peptide therapeutics for adult intestinal stem cell tumors. Dis Model Mech 2020;13

37. Brenner JC, Ateeq B, Li Y, Yocum AK, Cao Q, Asangani IA, et al. Mechanistic rationale for inhibition of poly(ADP-ribose) polymerase in ETS gene fusion-positive prostate cancer. Cancer Cell 2011;19:664–78

38. Stopsack KH, Su XA, Vaselkiv JB, Graff RE, Ebot EM, Pettersson A, et al. Transcriptomes of Prostate Cancer with TMPRSS2:ERG and Other ETS Fusions. Mol Cancer Res 2023;21:14–23

39. Meisel Sharon S, Pozniak Y, Geiger T, Werner H. TMPRSS2-ERG fusion protein regulates insulin-like growth factor-1 receptor (IGF1R) gene expression in prostate cancer: involvement of transcription factor Sp1. Oncotarget 2016;7:51375–92

40. Pettersson A, Lis RT, Meisner A, Flavin R, Stack EC, Fiorentino M, et al. Modification of the association between obesity and lethal prostate cancer by TMPRSS2:ERG. J Natl Cancer Inst 2013;105:1881–90

41. Mytsyk Y, Nakonechnyi Y, Dosenko V, Kowal P, Pietrus M, Gazdikova K, et al. The performance and limitations of PCA3, TMPRSS2:ERG, HOXC6 and DLX1 urinary markers combined in the improvement of prostate cancer diagnostics. Clin Biochem 2023;116:120–7

42. Jiang X, Liu Z, Wang H, Wang L. Discovery of lncRNA-Based ProsRISK Score in Serum as Potential Biomarkers for Improved Accuracy of Prostate Cancer Detection. J Cell Mol Med 2025;29:e70555

43. Rahim S, Beauchamp EM, Kong Y, Brown ML, Toretsky JA, Uren A. YK-4-279 inhibits ERG and ETV1 mediated prostate cancer cell invasion. PLoS One 2011;6:e19343

44. Rahim S, Minas T, Hong SH, Justvig S, Celik H, Kont YS, et al. A small molecule inhibitor of ETV1, YK-4-279, prevents prostate cancer growth and metastasis in a mouse xenograft model. PLoS One 2014;9:e114260

45. Winters B, Brown L, Coleman I, Nguyen H, Minas TZ, Kollath L, et al. Inhibition of ERG Activity in Patient-derived Prostate Cancer Xenografts by YK-4-279. Anticancer Res 2017;37:3385–96

46. Nhili R, Peixoto P, Depauw S, Flajollet S, Dezitter X, Munde MM, et al. Targeting the DNA-binding activity of the human ERG transcription factor using new heterocyclic dithiophene diamidines. Nucleic Acids Res 2013;41:125–38

47. Wang X, Liu W, Yang J, Tien JC, Chang Y, Mannan R, et al. A ligandable PNT domain establishes ERG as a directly targetable oncogenic driver in prostate cancer. Proc Natl Acad Sci U S A 2026;123:e2537437123

