## Supplementary figure S1 for "Targeting *TMPRSS2-ERG* Fusion-Driven Prostate Cancer using Novel HBS-α-Helicomimics"

1) 362 → TYR → 1<sup>st</sup> Residue in α-helix

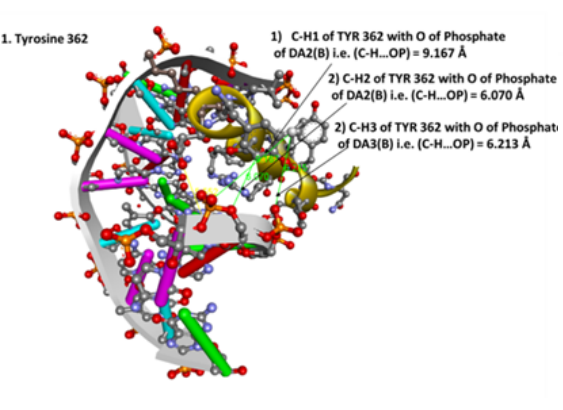

2) 363 → ASP → 2<sup>nd</sup> Residue in α-helix

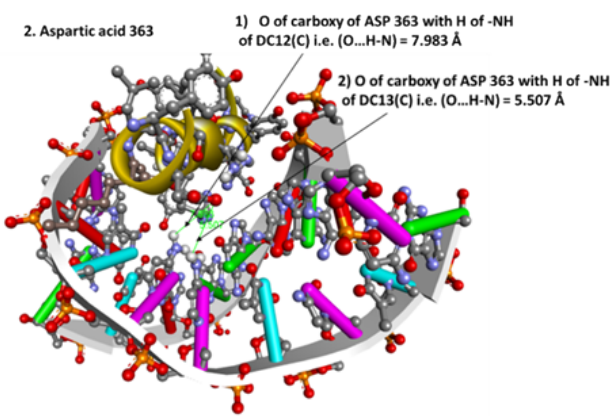

3) 364 → LYS → 3<sup>rd</sup> Residue in α-helix

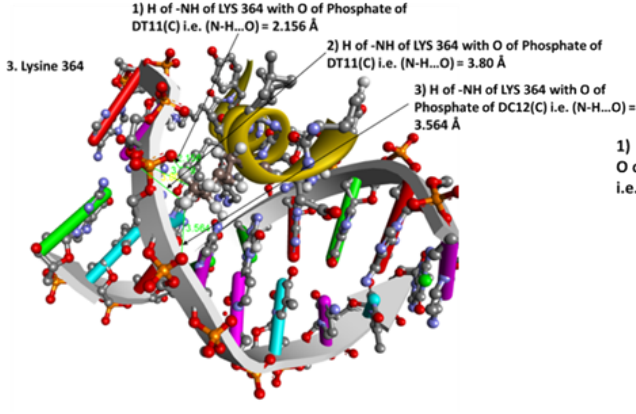

4) 365 → LEU → 4<sup>th</sup> Residue in α-helix

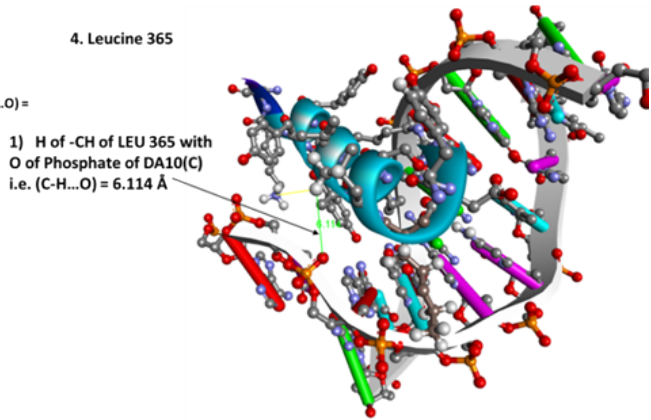

5) 366 → SER → 5<sup>th</sup> Residue in α-helix

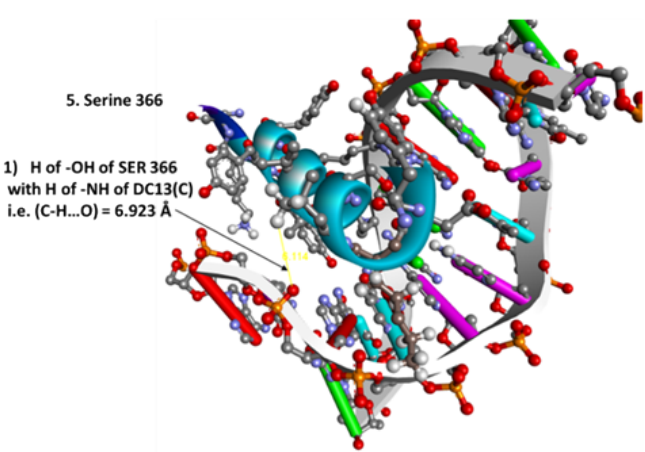

6) 367 → ARG → 6<sup>th</sup> Residue in α-helix

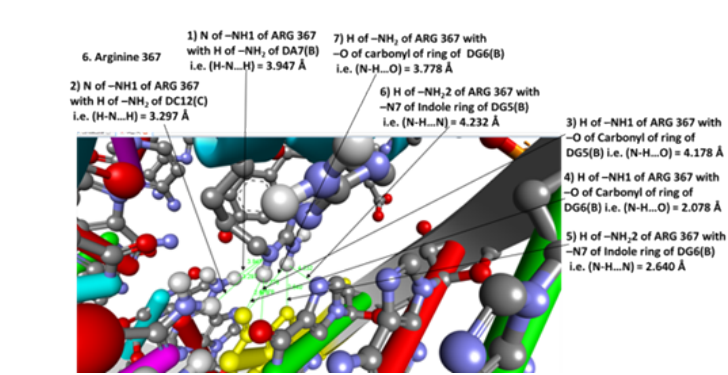

7) 368 → ALA → 7<sup>th</sup> Residue in α-helix

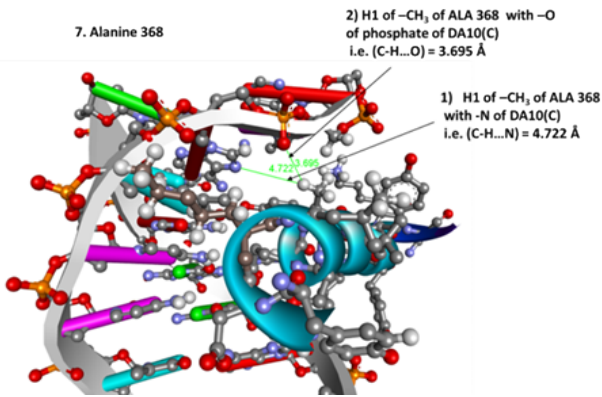

8) 369 → LEU → 8<sup>th</sup> Residue in α-helix

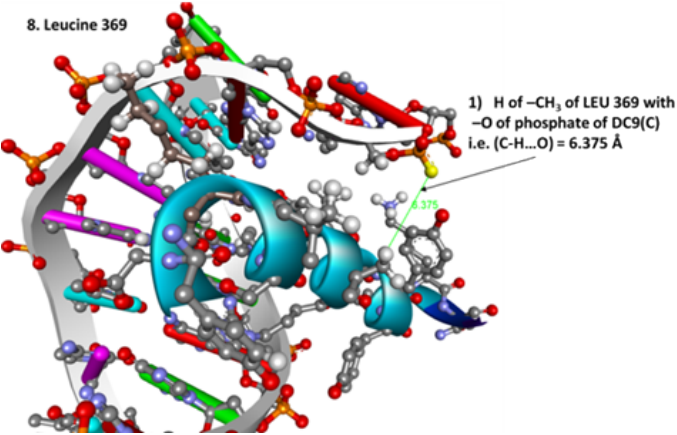

9) 370 → ARG → 9<sup>th</sup> Residue in α-helix

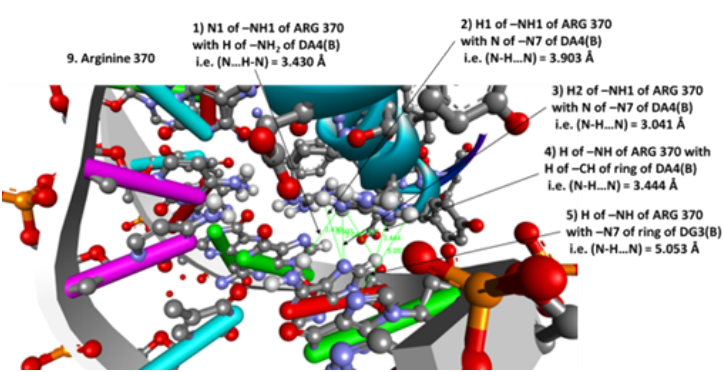

10) 371 → TYR → 10<sup>th</sup> Residue in α-helix

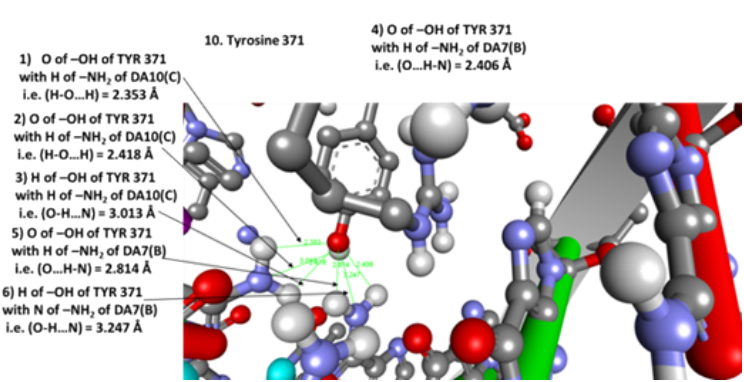

11) 372 → TYR → 11<sup>th</sup> Residue in α-helix

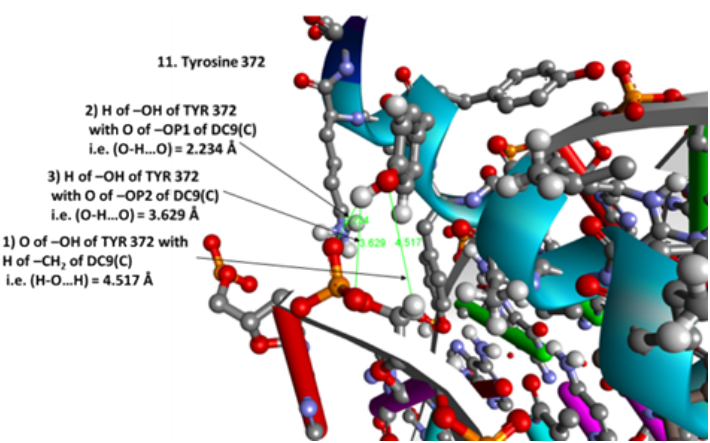

12) 373 → TYR → 12<sup>th</sup> Residue in α-helix

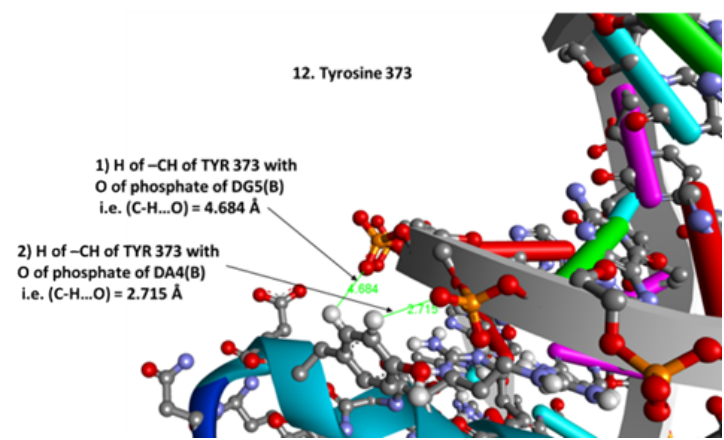

13) 375 → LYS → 14<sup>th</sup> Residue in α-helix

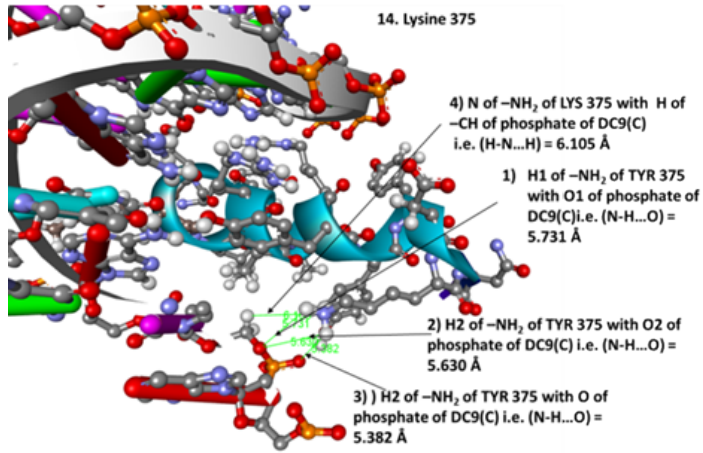

**Supplementary Figure S1:** Intermolecular contact profiles between ERG Helix 4 residues and dsDNA. 3D visual representations generated in BIOVIA Discovery Studio illustrating specific electrostatic, hydrogen-bonding, and van der Waals interactions between individual amino acid side chains within the recognition sequence <sup>362</sup>YDKLSRALRYYYDK<sup>375</sup>N and the corresponding nucleobases or phosphodiester backbone of the target DNA duplex.
