## Supplementary figure S2 for "Targeting *TMPRSS2-ERG* Fusion-Driven Prostate Cancer using Novel HBS-α-Helicomimics"

**Scheme 1.** Synthesis of HBS-constrained single helical turn peptide mimic:

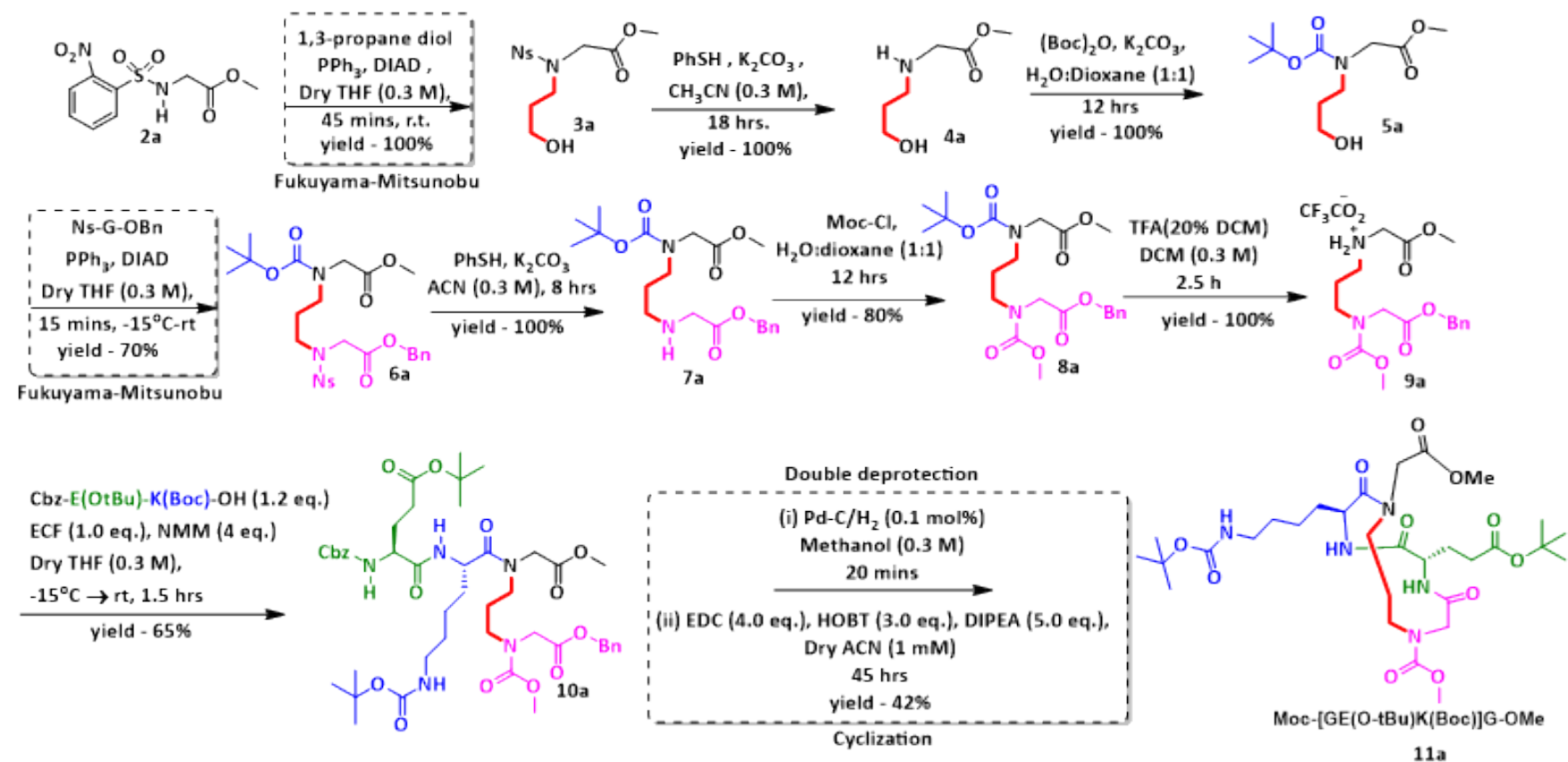

**Scheme 2.** Synthesis of HBS-constrained extended helicomic:

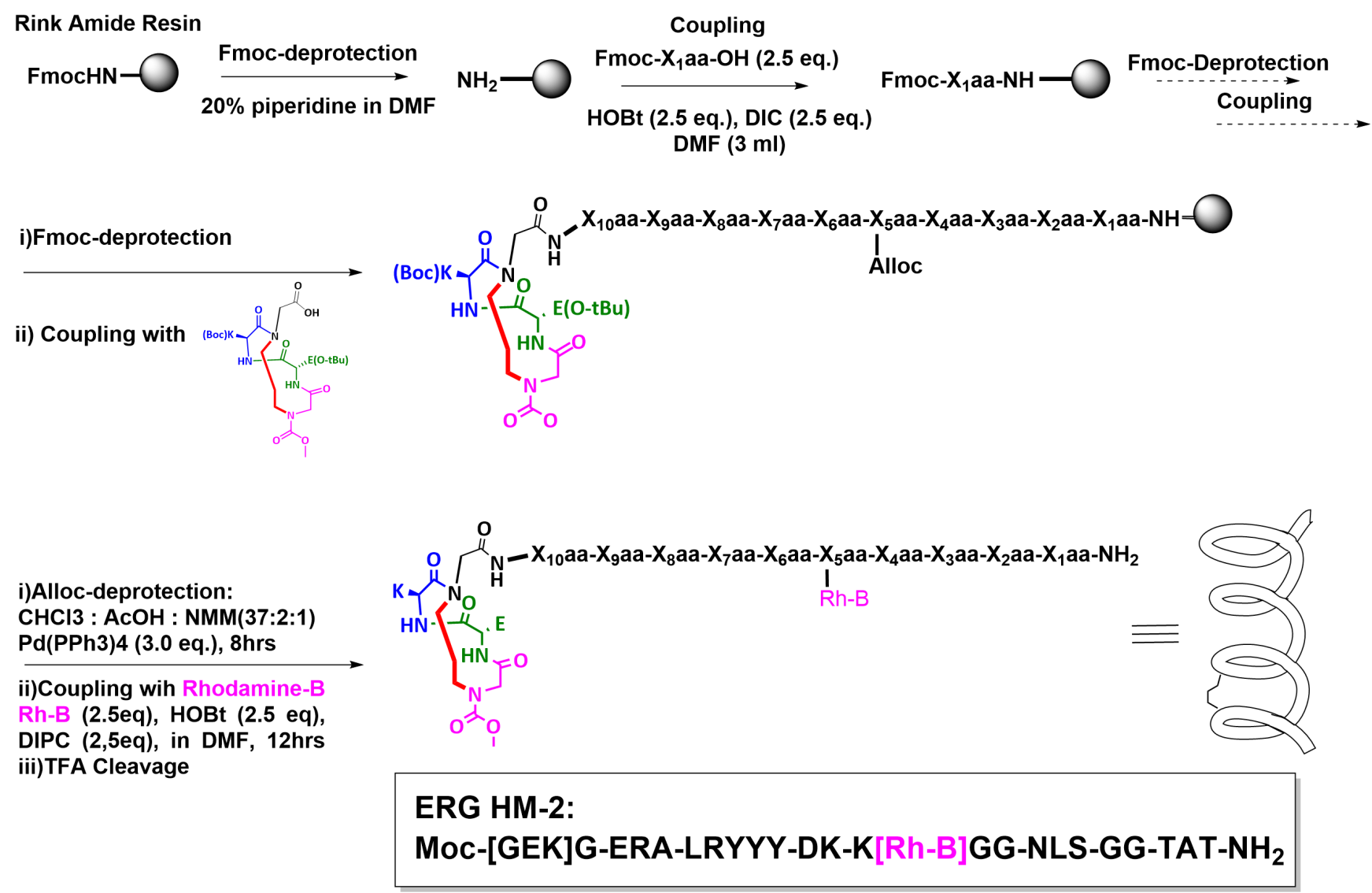
