## Supplementary figure S3 for "Targeting *TMPRSS2-ERG* Fusion-Driven Prostate Cancer using Novel HBS-α-Helicomimics"

A

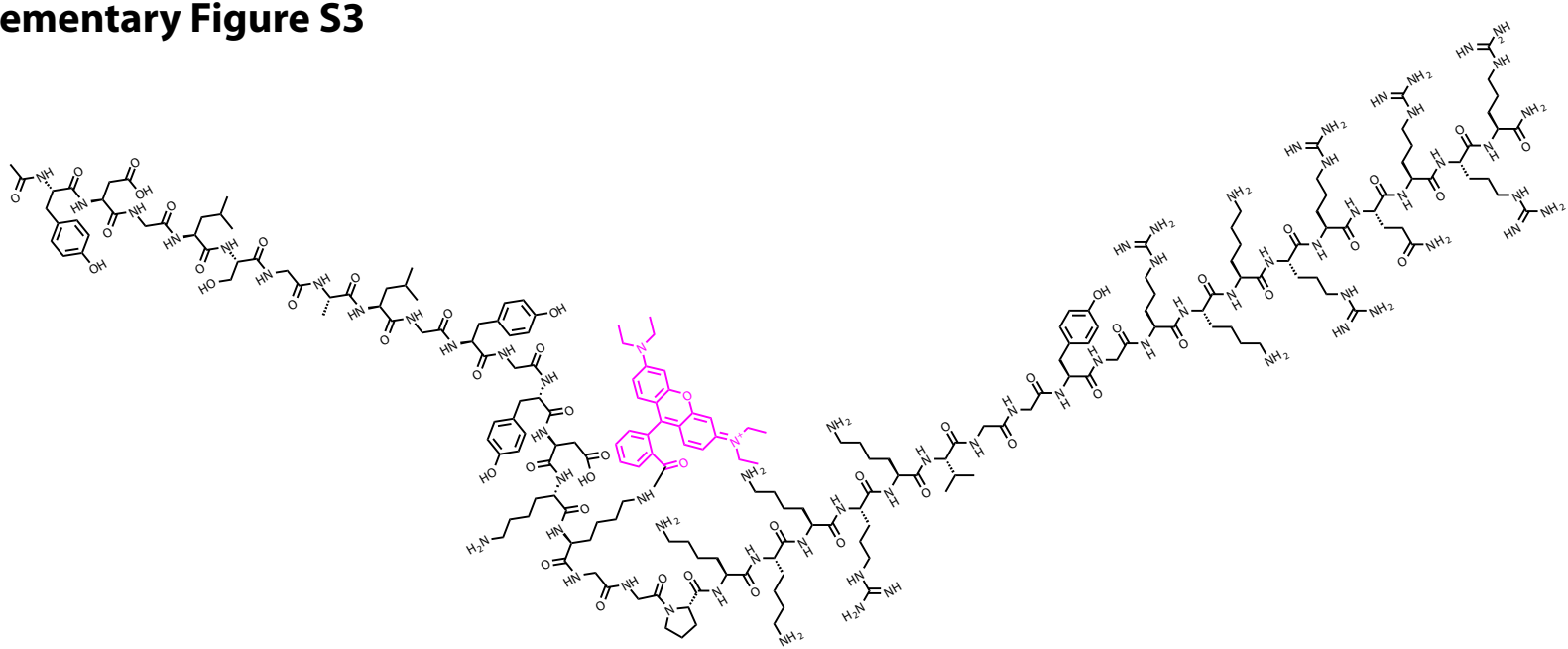

Control peptide: Ac-YDGL-SGA-LGYGY-DK-K(Rh-B)-GG-PKKKRKV-GG-YGRKKRRQRRR-NH<sub>2</sub>

B

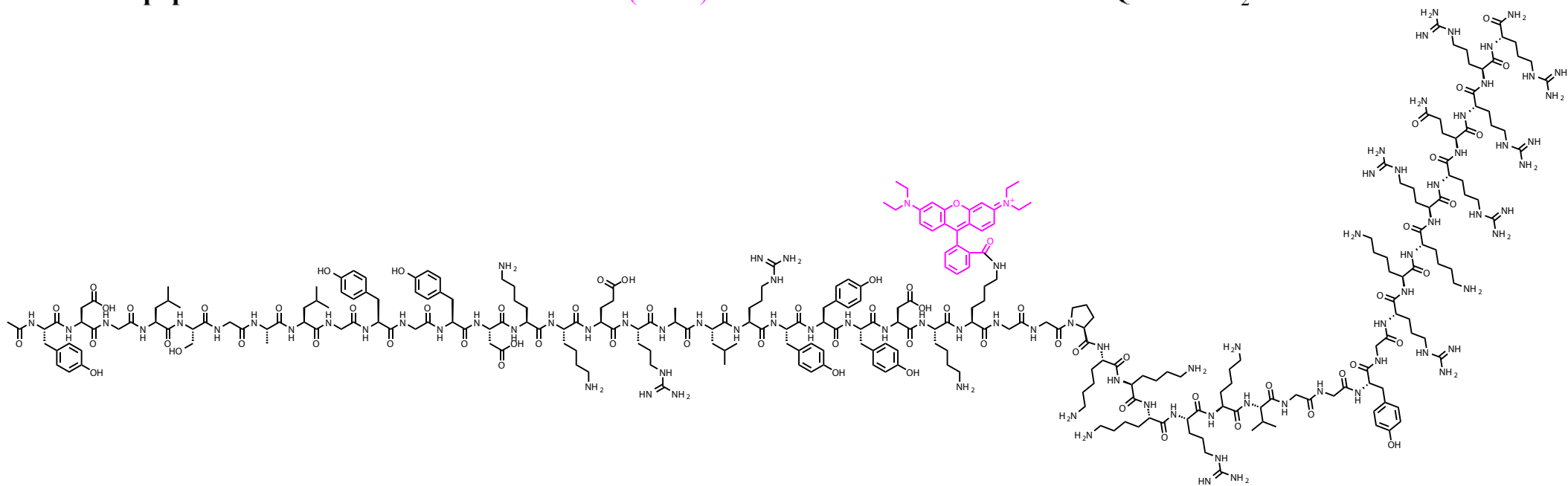

ERG HM-1: Ac-KD-YYYRL-ARS-LKDY-K(Rh-B)-GG-PKKKRKV-GG-YGRKKRRQRRR-NH<sub>2</sub>

C

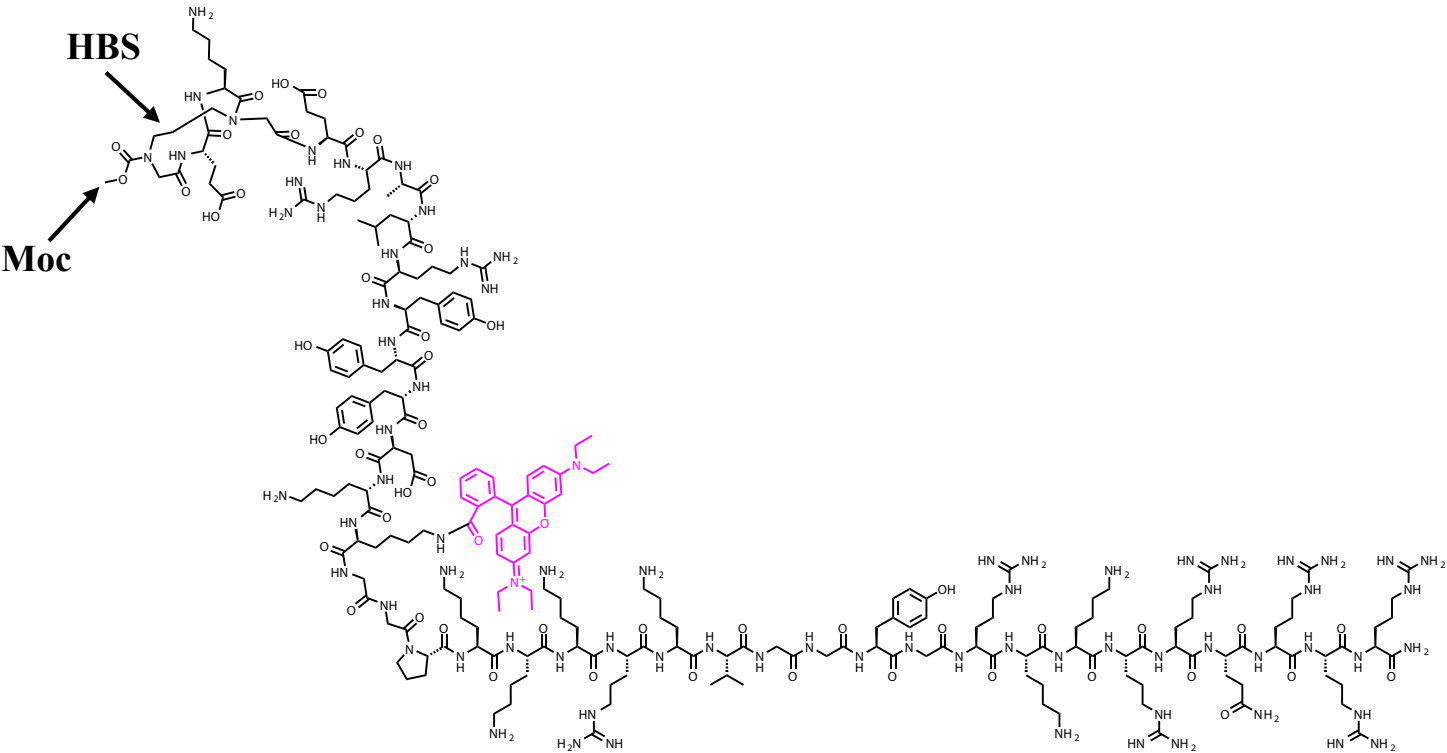

ERG HM-2: Moc-[GEK]G-ERA-LRYYYDK-K(Rh-B)-GG-PKKKRKV-GG-YGRKKRRQRRR-NH<sub>2</sub>.

Supplementary Figure S3: Chemical structures of synthesised A) Control peptide B) ERG HM-1 and C) ERG HM-2.
