## Supplementary Methods for "Targeting *TMPRSS2-ERG* Fusion-Driven Prostate Cancer using Novel HBS-α-Helicomimics"

##### \*Corresponding Author:

### Supplementary Methods:

#### Structural Analysis and Mapping of Protein-DNA Contacts

To establish the structural basis for helicomimic design, the native co-crystal structure of the ERG transcription factor bound to dsDNA (PDB: 6VGG) was analyzed. Non-covalent contacts between the recognition Helix 4 (H4) and the cognate major-groove binding tracks were mapped using BIOVIA Discovery Studio. Specific non-covalent interactions—encompassing hydrogen bonding, salt bridges, electrostatic contacts, and van der Waals interactions—were identified based on defined donor-acceptor distance thresholds (1,2): $\text{P-O} \cdots \text{H} \leq 4.0 \text{ \AA}$ (phosphate backbone interactions), $(\text{N/O})\text{-X} \cdots \text{H-Y} \leq 3.8 \text{ \AA}$ (standard hydrogen bonds), $(\text{N/O})\text{-X} \cdots \text{H-C} \leq 3.2 \text{ \AA}$ (weak hydrogen bonding/dipole interactions), $\text{C} \cdots \text{H-C} \leq 3.2 \text{ \AA}$ (steric and van der Waals contacts). An angular criterion ranging from $180^\circ$ (ideal) down to a $135^\circ$ cutoff was applied to confirm geometric directional validity. The individual interaction diagrams depicting spatial proximity and specific residue-to-base contacts are shown below (**Fig. S1**).

#### General Solid-Phase Peptide Synthesis (SPPS) and On-Resin Rhodamine-B Functionalization

Peptide sequences were assembled manually via Fmoc-based solid-phase peptide synthesis on Rink Amide resin (loading: 0.661 mmol/g) to provide a C-terminal primary carboxamide upon cleavage. The resin was pre-swollen in anhydrous DCM (3 x 10 min) followed by anhydrous DMF (3 x 10 min) under gentle agitation. Iterative removal of the N-alpha-Fmoc group was carried out using 20% piperidine in DMF (v/v, 2 x 15 min), followed by systematic washing with DMF (5 x 2 min), DCM (3 x 2 min), and DMF (3 x 2 min). Individual amino acid couplings were performed sequentially from the C- to N-terminus using standard Fmoc-protected L-amino acids bearing acid-labile side-chain protections [Boc for Lys, Pbf for Arg, tBu for Tyr and Ser, OtBu for Glu and Asp, and Trt for Gln]. The amino acid derivative (2.5 equiv relative to resin loading) was pre-activated with HOBt (2.5 equiv) and DIPC (2.5 equiv) in anhydrous DMF (3 mL) for 5 min, transferred to the peptidyl-resin, and agitated for 1.5–2 h at room temperature.

To introduce the fluorophore handle at a defined position, an orthogonally protected lysine residue, Fmoc-Lys(Alloc)-OH, was incorporated into the sequence during backbone assembly. Following complete elongation of the peptide chain and coupling of the N-terminal HBS macrocyclic building block, selective on-resin deprotection of the allyloxycarbonyl (Alloc) group was performed. The peptidyl-resin was swollen in anhydrous $\text{CHCl}_3$ and treated with a deprotection cocktail containing $\text{CHCl}_3$, $\text{AcOH}$, and $\text{NMM}$ in a 37:2:1 (v/v/v) ratio, followed by the addition of $\text{Pd(PPh}_3)_4$ (3.0 equiv) and vortexing for 8 hours. The resin was then filtered and washed with $\text{CHCl}_3$ (5 x 2 min), $\text{DMF}$ (5 x 2 min), $\text{DCM}$ (3 x 2 min), and $\text{DMF}$ (3 x 2 min).

Direct fluorophore conjugation was carried out by treating the resin with a pre-activated solution of Rhodamine B (2.5 equiv), $\text{HOBt}$ (2.5 equiv), and $\text{DIC}$ (2.5 equiv) in anhydrous $\text{DMF}$ (3 mL). The reaction vessel was shielded from light and agitated for 12 h at room temperature. The resin was washed thoroughly with $\text{DMF}$ (5 x 2 min), $\text{DCM}$ (5 x 2 min), and $\text{MeOH}$ (3 x 2 min) until the wash filtrates were colorless, then dried under vacuum prior to global acidolytic cleavage and side-chain deprotection (**Fig. S2**).

Using the synthetic schemes above, the following peptides were synthesized (**Fig. S3**):

**Control peptide:** $\text{Ac-YDGL-SGA-LGYGY-DK-K(Rh-B)-GG-PKKKRKV-GG-YGRKKRRQRRR-NH}_2$

**ERG HM-1:** $\text{Ac-KD-YYYRL-ARS-LKDY-K(Rh-B)-GG-PKKKRKV-GG-YGRKKRRQRRR-NH}_2$.

**ERG HM-2:** $\text{Moc-[GEK]G-ERA-LRYYYDK-K(Rh-B)-GG-PKKKRKV-GG-YGRKKRRQRRR-NH}_2$.

##### Synthesis, HRMS, LC-MS and HPLC of peptides:

**Synthesis of Cbz-Glu(O-tBu)-OH ((S)-2-(((benzyloxy)carbonyl)amino)-5-(tert-butoxy)-5-oxopentanoic acid) (a):**

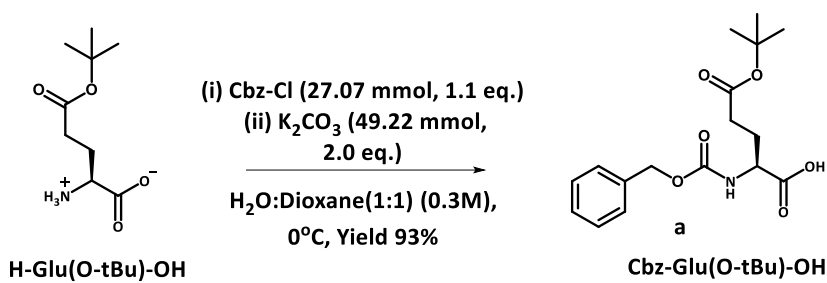

To a cold ($0^\circ\text{C}$) stirring mixture of side chain tert-butyl protected glutamic acid (5.0g, 24.61 mmol) and $\text{K}_2\text{CO}_3$ (6.79 g, 49.22 mmol) in water (41.0 ml) was added a solution of benzyl chloroformate (3.87 ml, 27.07 mmol) in dioxane (41.0 ml) dropwise. The mixture was stirred at $0^\circ\text{C}$ for 1 h and warmed to stir at ambient conditions until TLC indicated the complete consumption of starting material (10 h). The dioxane was removed under vacuum to get an aqueous solution which was acidified with 1N HCl solution and extracted with EtOAc (3 X 20.0 ml). The organic layer was dried over anhyd. Na<sub>2</sub>SO<sub>4</sub> and the solvent was removed under vacuum to get a hygroscopic white solid (7.7 g, 93 %), which was used for further reaction without any purification.

**Synthesis of Boc-Glu(O-tBu)-Lys(Boc)-OMe (methyl N<sup>2</sup>((S)2(((benzyloxy)carbonyl)amino)-5-(tert-butoxy)-5-oxopentanoyl)-N<sup>6</sup>-(tert-butoxycarbonyl)-L-lysinate) (b):**

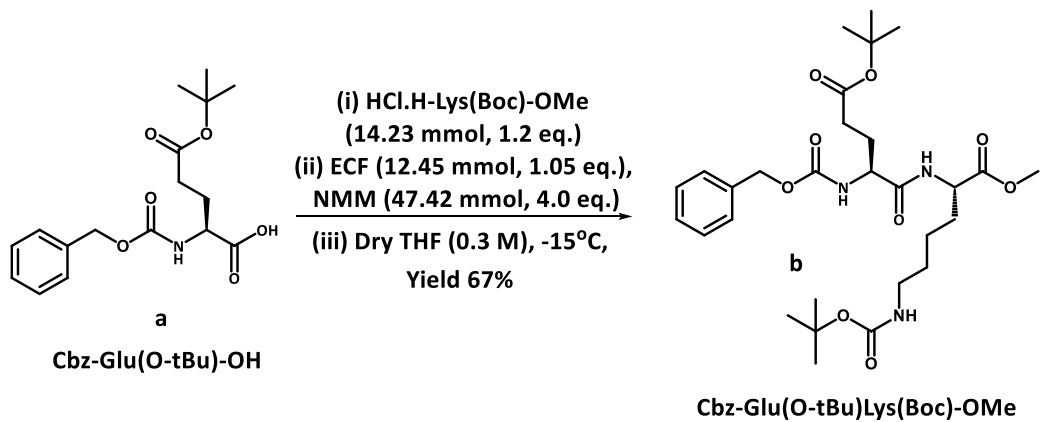

To a cold (-15°C) stirring solution of Cbz-Glu(O-tBu)-OH (**a**) (4.0 g, 11.86 mmol) in dry THF (25 ml) was added NMM (1.94 ml, 17.78 mmol) and ECF (1.19 ml, 12.45 mmol) and stirred until TLC indicated that all the acid had been consumed to form corresponding mixed anhydride (4 min). To this was added a solution of HCl.H-Lys(Boc)-OMe (3.70 g, 14.23 mmol) in dry THF (15.0 ml) and NMM (3.23 ml, 29.64 mmol). After 2 h the ice-salt bath was removed and the mixture was stirred at ambient conditions for further 5 h. Removal of solvent resulted in a residue which was dissolved in ethyl acetate (EtOAc) (60.0 ml), washed with water (2 X 25.0 ml), 1N HCl (2 X 25.0 ml) and saturated NaHCO<sub>3</sub> (2 X 25.0ml). The organic layer was dried over anhyd. Na<sub>2</sub>SO<sub>4</sub> and concentrated to get a residue which was subjected to purification by silica gel flash column chromatography (EtOAc : Hexane – 2:1) to yield the desired product as a white solid (4.6 g, 7.94 mmol, 67% yield); (TLC- EtOAc:Hexane (1:1) – $R_f$ = 0.6); <sup>1</sup>H NMR (400 MHz, CDCl<sub>3</sub>, 10 mM) δ ppm: 7.37 (s, 1H, H<sup>Aro</sup>Cbz), 7.36 (s, 4H, H<sup>Aro</sup>Cbz), 6.96 (s, 1H, H<sup>N</sup>Lys), 5.71 (s, 1H, H<sup>N</sup>Glu), 5.13 (s, 2H, H<sup>CH2</sup>Cbz) & 4.71 (s, 1H, H<sup>N side chain</sup>Lys), 4.57 (d, J = 5.001 Hz, 1H, H<sup>α</sup>Lys) & 4.28 (q, J = 6.78 Hz, 1H, H<sup>α</sup>Glu), 3.75 (s, 3H, H<sup>Me</sup>OMe), 3.097 (d, J = 6.05 Hz, 2H, H<sup>ω</sup>Lys), 2.49 – 2.36 (m, 2H, H<sup>γ</sup>Glu), 2.146 – 2.06 (m, 1H, H<sup>β</sup>Glu) and 1.997 – 1.92 (m, 1H, H<sup>β</sup>Glu), 1.904 – 1.83 (m, 1H, H<sup>β</sup>Lys), 1.75 – 1.699 (m, 3H, H<sup>β</sup>Lys & H<sup>δ</sup>Lys), 1.465 (s, 9H, H<sup>Me</sup>t-Bu), 1.45 (s, 9H, H<sup>Me</sup>Boc), 1.38 – 1.29 (m, 2H, H<sup>γ</sup>Lys); <sup>13</sup>C NMR (100 MHz, CDCl<sub>3</sub>, 60 mM) δ ppm: 172.6, 172.2, 171, 156, 155.8, 136, 128.3, 127.8, 80.8, 78.9, 66.7, 53.8, 52.1, 39.8, 31.3, 29, 27.8, 22.1; FT-IR ($\tilde{\nu}$ cm<sup>-1</sup>):

3912.8, 3861.7, 3804.1, 3715.3, 3661, 3581, 3533, 3320, 3052.4, 2937.3, 2771.7, 2635, 2507, 2418, 2300, 1987.3, 1867, 1822, 1691.4, 1522, 1443, 1366, 1245.2, 1146, 1035, 971, 821, 732, 668; **HRMS** $m/z$ Calculated for $C_{12}H_{22}N_2O_5Na$ 602.3053, Found 602.3055.

Synthesis of Ns-Gly-OMe (methyl 2-(2-nitrophenylsulfonamido)acetate) (**2a**):

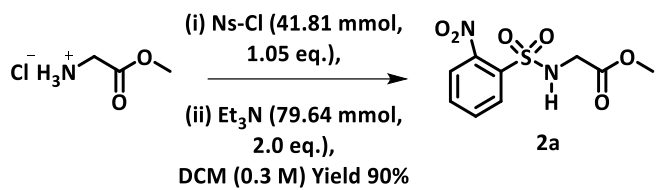

To a cold (0°C) stirring mixture of methylglycinate hydrochloride (5.0 g, 39.82 mmol) and o-nitrobenzenesulfonyl chloride (NsCl) (9.26 g, 41.81 mmol) in DCM (133.0 ml) was added triethylamine (TEA) (11.12 ml, 79.64 mmol) drop-wise and the mixture was stirred at room temperature until TLC indicated the complete consumption of the starting material. Removal of solvent resulted in a residue which was dissolved in EtOAc (70.0 ml), washed with water (2 x 20.0 ml) and 1N HCl solution (2 X 20.0 ml). The organic layer was dried over anhyd. Na<sub>2</sub>SO<sub>4</sub> and concentrated to get a residue which was subjected to purification by silica gel flash column chromatography (EtOAc:Hexane -2:3) to yield the desired product (Ns-Gly-OMe, **1a**) as a white solid (9.82 g, 35.84 mmol, 90% yield); m.p. 110°C; (TLC- EtOAc:Hexane (1:1) – $R_f$ = 0.5); **<sup>1</sup>H NMR** (400 MHz, CDCl<sub>3</sub>, 10 mM) $\delta$ ppm: 8.11-8.08 (m, 1H, H<sup>Aro</sup>Ns), 7.95-7.93 (m, 1H, H<sup>Aro</sup>Ns), 7.76-7.73 (m, 2H, H<sup>Aro</sup>Ns), 6.03 (s, 1H, H<sup>N</sup>Gly), 4.02 (d, J = 5.86 Hz, 2H, H<sup>α</sup>Gly), 3.61 (s, 3H, H<sup>Me</sup>OMe); **<sup>13</sup>C NMR** (100 MHz, CDCl<sub>3</sub>, 60 mM) $\delta$ ppm: 169.2, 148.0, 134.2, 133.9, 133.1, 130.8, 125.8, 52.7, 45.0; **FT-IR** ($\tilde{\nu}$ cm<sup>-1</sup>): 3324, 3099, 1754, 1639, 1593, 1537, 1465, 1417, 1358, 1335, 1257, 1239, 1160, 1126, 1056, 1008; **HRMS** $m/z$ Calculated for $C_9H_{10}N_2O_6SNa$ 297.0157, Found 297.0161.

#### Synthesis of methyl 2-((tert-butyloxycarbonyl)(3-hydroxypropyl)amino) acetate (4a):

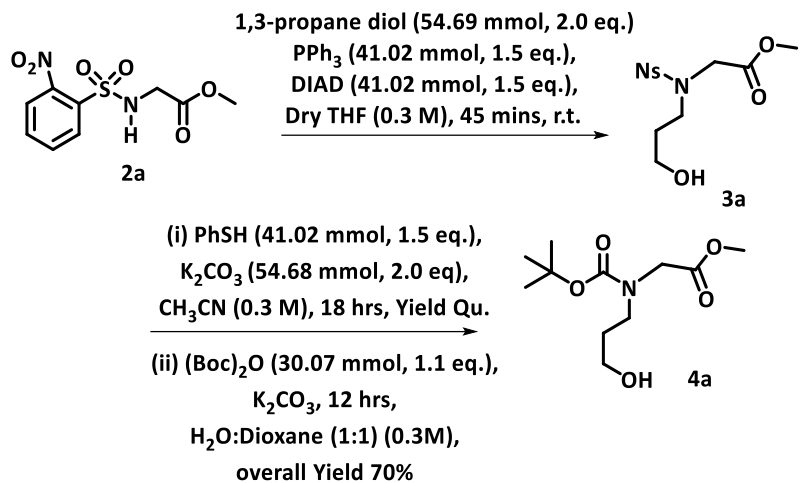

To a cold (0°C) stirring mixture of Ns-Gly-OMe (**1a**) (7.5 g, 27.34 mmol) and triphenylphosphine (PPh<sub>3</sub>) (10.74 g, 41.02 mmol) in dry THF (91.0 ml) under N<sub>2</sub> atmosphere was added 1,3-propanediol (3.95 ml, 54.69 mmol) drop-wise followed by diisopropyl azodicarboxylate (DIAD) (8.12 ml, 41.02 mmol). After 15 mins of stirring the ice bath was removed and the reaction was continued at ambient conditions until TLC indicated the complete consumption of the starting material. Removal of the solvent resulted in a residue that was dissolved in EtOAc (50.0 ml) and washed with water (2 X 25.0 ml). The organic layer was concentrated to get a residue which was immediately subjected to denosylation reaction conditions by the addition of thiophenol (4.2 ml, 41.02 mmol) and K<sub>2</sub>CO<sub>3</sub> (7.54 g, 54.68 mmol) in acetonitrile (91.0 ml). The reaction was continued at ambient conditions until TLC indicated the complete consumption of the starting material. After that, the solvent was removed and the resulting residue was acidified using 1N HCl (50.0 ml). The aqueous mixture was thoroughly washed with diethyl ether (3 x 20.0 ml) to remove all the water-insoluble organic impurities.

The resulting aqueous solution was then treated with K<sub>2</sub>CO<sub>3</sub> until it reached a pH of 10. and di-tert-butylidicarbonate (6.90 ml, 30.07 mmol) was added and the reaction mixture was stirred at ambient conditions until TLC indicated the complete consumption of the starting material. Removal of the solvent resulted in a residue which was dissolved in EtOAc (70.0 ml) and the organic layer was washed with water (3 x 20 mL) and dried over anhyd. Na<sub>2</sub>SO<sub>4</sub> and concentrated to get a residue which was subjected to purification by silica gel flash column chromatography (EtOAc:Hexane -1:10) to yield the desired product as a viscous oil (4.73g, 19.12 mmol, 70%.

yield); (TLC- EtOAc:Hexane (1:0) – $R_f$ = 0.5); $^1\text{H NMR}$ (400 MHz, $\text{CDCl}_3$, 10 mM) $\delta$ ppm: (mixture of two rotamers) 3.86 (s) & 3.96 (s) (2H, $\text{H}^{\text{a}}\text{Gly}$), 3.74 (s, 3H, $\text{H}^{\text{Me}}\text{OMe}$), 3.65, (t, $J$ = 5.3 Hz), & 3.71-3.68 (m), (2H, $\text{HPrp}^{\text{c}}$); 3.45 (t, $J$ = 6.0 Hz) & 3.41-3.38 (m), (2H, $\text{HPrp}^{\text{a}}$); 1.66-1.64 (m) & 1.77-1.74 (m) (2H, $\text{HPrp}^{\text{b}}$); 1.48 (s) & 1.42 (s), (9H, $\text{H}^{\text{Me}}\text{Boc}$); $^{13}\text{C NMR}$ (100 MHz, $\text{CDCl}_3$, 60 mM) $\delta$ ppm: (mixture of two rotamers) 170.8, 170.6, 156.7, 155.9, 81.2, 80.9, 60.0, 58.6, 52.2, 49.8, 49.2, 45.5, 44.8, 31.4, 30.8, 28.5, 28.3; **FT-IR** ($\tilde{\nu}$ $\text{cm}^{-1}$): 3413, 2974, 2956, 2875, 1753, 1682, 1479, 1437, 1406, 1368, 1313, 1250, 1210, 1061; HRMS $m/z$ Calcd for $\text{C}_{11}\text{H}_{21}\text{NO}_5\text{Na}$ 270.1317, Found 270.1320.

**Synthesis of (S)-benzyl 2-((3-((tert-butyloxycarbonyl)(2-methoxy-2-oxoethyl) amino) propyl) (methoxycarbonyl)amino)propanoate (7a):**

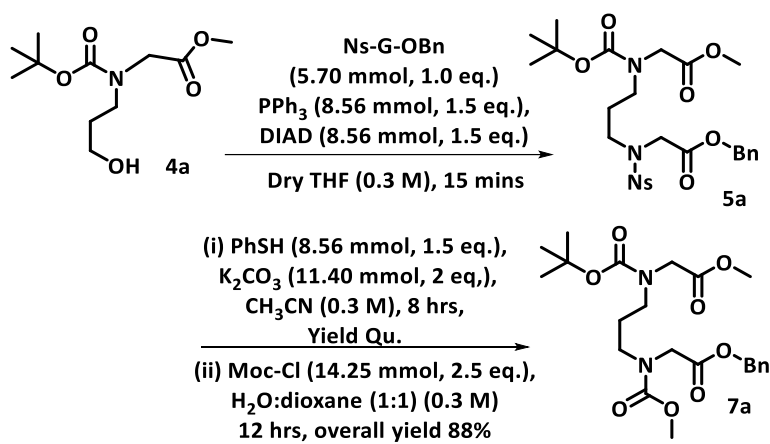

To a cold ($0^\circ\text{C}$) stirring mixture of $\text{Ns-Gly-OBn}$ (2.0 g, 5.70 mmol) and $\text{PPh}_3$ (2.24 g, 8.56 mmol) in dry THF (19.0 ml) under $\text{N}_2$ atmosphere was added $\text{N-Boc-(3-hydroxypropyl) benzylglycinate}$ (**4a**) (2.11 gm, 8.56 mmol) dissolved in THF followed by $\text{DIAD}$ (1.69 ml, 8.56 mmol). After 15 minutes the ice bath was removed, and the mixture was continued to stir at ambient conditions until TLC indicated the complete consumption of the starting material. Removal of the solvent resulted in a residue which was dissolved in EtOAc (30.0 ml) and washed with water (2 X 15.0 ml). The organic layer was dried over anhyd. $\text{Na}_2\text{SO}_4$ and concentrated to get a residue which was subjected to denosylation reaction by the addition of thiophenol (0.87 ml, 8.56 mmol) and $\text{K}_2\text{CO}_3$ (1.57g, 11.40 mmol) in acetonitrile (19.0 ml). The mixture was continued to stir at ambient conditions until TLC indicated the complete consumption of the starting material. The solvent was removed followed by acidification using 1N $\text{HCl}$ (20.0 ml). The aqueous mixture was washed with diethyl ether to remove all the water-insoluble organic impurities.

The aqueous solution was then treated with $K_2CO_3$ until it reached a pH of 10. This was followed by the addition of methylchloroformate (MCF) (1.10 ml, 14.25 mmol) to the reaction mixture. The mixture was continued to stir at ambient conditions until TLC indicated the complete consumption of the starting material. Removal of solvent resulted in a residue which was dissolved in EtOAc (30.0 ml), washed with water (2 X 10.0 ml) and the organic layer was dried over anhyd. $Na_2SO_4$ and concentrated to get a residue which was subjected to purification by silica gel flash column chromatography (EtOAc:Hexane -1:5) to yield the desired product as a viscous oil (2.26g, 4.99 mmol, 88% yield); (TLC- EtOAc:Hexane (1:1) – $R_f$ = 0.4); **$^1H$ NMR** (400 MHz, $CDCl_3$, 10 mM) $\delta$ ppm: (mixture of four rotamers) 7.40-7.30 (m, 5H, $H^{AroBn}$), 5.16 (s, 2H, $H^{CH_2Bn}$), 4.055 (d, $J$ = 4.08 Hz, 1H, $H^{\alpha}Gly$), 3.99 (d, $J$ = 11.65 Hz, 1H, $H^{\alpha}Gly$), 3.93 (s, 1H, $H^{\alpha}Gly$), 3.84 (d, $J$ = 13.15, 1H, $H^{\alpha}Gly$), 3.75 – 3.70 (m), 3.64 – 3.59 (m) ($H^{MeOMe}$ & $H^{MeMoc}$), 3.36 – 3.25 (m, 4H, $H^{Prp^c}$ & $H^{Prp^a}$), 1.77 – 1.71 (m, 2H, $H^{Prp^b}$), 1.45 (s) & 1.40 (s) (9H, $H^{MeBoc}$); **$^{13}C$ NMR** (100 MHz, $CDCl_3$, 60 mM) $\delta$ ppm: (mixture of four conformers) 170.8, 170.7, 170.6, 169.9, 169.7, 157.1, 156.6, 155.7, 155.2, 135.5, 135.4, 128.6, 128.5, 80.52, 80.46, 80.37, 80.26, 66.94, 66.86, 66.81, 53.0, 52.8, 52.0, 49.4, 49.3, 49.2, 49.0, 48.9, 46.8, 46.7, 46.4, 46.3, 45.9, 28.34, 28.20, 27.5, 27.2, 27.1, 26.7, ; **FT-IR** ($\tilde{\nu} \text{ cm}^{-1}$): 3583, 2975, 2954, 1752, 1703, 1476, 1403, 1367, 1245, 1209, 1175, 1143, 1070, 1012; **HRMS** $m/z$ Calculated for $C_{23}H_{34}N_2O_8K$ 491.1796, Found 491.1796.

#### Synthesis of precursor of cyclisation (9a):

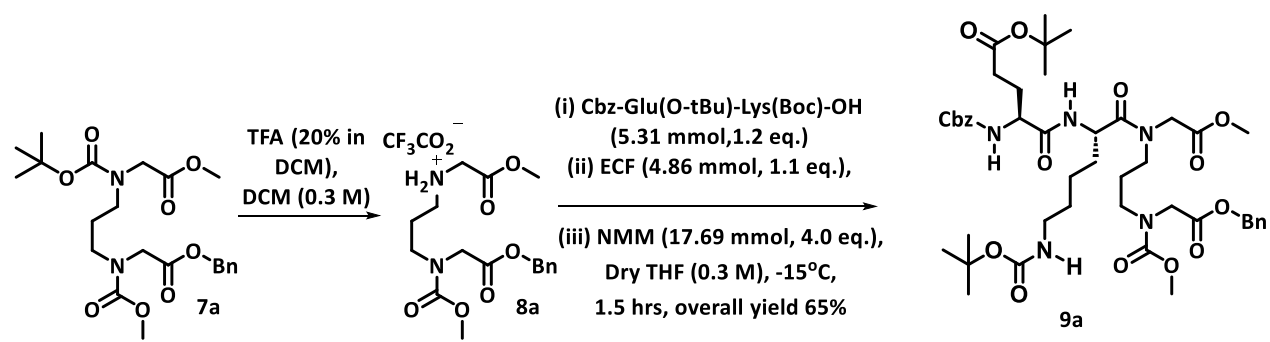

To a cold, ($0^\circ\text{C}$) solution of **7a** (2.0 g, 4.42 mmol) in DCM (11.79 ml) was added trifluoroacetic acid (TFA) (2.95 ml) dropwise and the mixture was allowed to stir for 2 hrs by which time TLC indicated the complete consumption of the starting material. Removal of the solvent resulted in the Boc-protected product **8a** as a liquid (2.06 g, 4.42 mmol, quantitative yield) which was used for further reaction as it was.

To a cold (-15°C) solution of Cbz-Glu(O-tBu)-Lys(Boc)-OH (3.00 g, 5.31 mmol) in dry THF (10.7 ml) and NMM (0.723 ml, 6.63 mmol) was added ECF (0.465 ml, 4.86 mmol). The mixture was stirred for 4 min when TLC indicated that all the acid was consumed to form the corresponding mixed anhydride. To this mixture was added a solution of **8a** (2.06 g, 4.42 mmol) in dry THF (4.0 ml) followed by NMM (1.21 ml, 11.06 mmol). After 2 h of vigorous stirring, the mixture was warmed and stirred until TLC indicated the complete consumption of starting material. Removal of solvent resulted in a residue which was dissolved in EtOAc (40.0 ml), washed with water (2 X 20.0 ml) and 1N HCl (2 X 20.0 ml) and saturated NaHCO<sub>3</sub> (2 X 20.0 ml). The organic layer was dried over anhyd. Na<sub>2</sub>SO<sub>4</sub> and concentrated to get a residue which was subjected to purification by silica gel flash column chromatography (EtOAc:Hexane – 6:1) to yield the desired product as a liquid (1.79g, 1.99 mmol, 45% yield); (TLC- EtOAc:Hexane (3:1) – *R<sub>f</sub>* = 0.35); **<sup>1</sup>H NMR** (400 MHz, CDCl<sub>3</sub>, 10 mM) δ ppm: (mixture of four rotamers) 8.13 – 8.02 (m, 1H, H<sup>N</sup>Lys), 7.34 – 7.33 (m, 10H, H<sup>Aro</sup>Bn, H<sup>Aro</sup>Cbz), 6.99 – 6.93 (m, 1H, H<sup>N</sup>Glu), 5.74 – 5.67 (m, 1H, H<sup>Nside chain</sup> Lys), 5.16 (s, 2H, H<sup>CH<sub>2</sub></sup>Bn), 5.097 – 5.084 (m, 2H, H<sup>CH<sub>2</sub></sup>Cbz), 4.865 (s, 1H, H<sup>α</sup>Lys), 4.346 – 4.195 (m, 2H, H<sup>α</sup>Gly1), 4.07 – 4.00 (m, 2H, H<sup>α</sup>Gly4), 3.96 (s, 1H, H<sup>α</sup>Glu), 3.79 – 3.74 (m, 3H, H<sup>Me</sup>Moc), 3.709 (d, *J* = 1.888, 3H, H<sup>Me</sup>OMe), 3.627 – 3.5988 (m, 2H, H<sup>ω</sup>Lys), 3.396 – 3.36 (m, 4H, HPrp<sup>c</sup> & HPrp<sup>a</sup>), 3.057 (bs, 2H, H<sup>β</sup>Glu), 2.3422 (bs, 2H, H<sup>γ</sup>Glu), 2.0616 – 2.015 (m, 2H, HPrp<sup>b</sup>), 1.911 – 1.86 (m, 2H, H<sup>β</sup>Lys), 1.759 – 1.743 (m, 2H, H<sup>δ</sup>Lys), 1.416 (s, 18H, H<sup>Me</sup>Boc + tBu), 1.246 (bs, 2H, H<sup>γ</sup>Lys); **<sup>13</sup>C NMR** (100 MHz, CDCl<sub>3</sub>) δ ppm: (mixtures of four conformers) 172.5, 172.1, 171, 169.7, 169.6, 169.3, 163.2, 156.7, 156, 136.1, 135.2, 135.1, 128.5, 128.45, 128.3, 128.2, 128.1, 128.08, 127.98, 127.92, 127.87, 127.6, 80.7, 79, 66.9, 54.2, 53.3, 53.02, 52.9, 52.7, 52.45, 52.13, 52.03, 49.4, 49.3, 49.2, 48.9, 48.6, 47.78, 47.6, 46.7, 46.4, 46.3, 46.01, 45.88, 44.08, 43.9, 40.08, 32.09, 31.74, 31.48, 29.5, 29.15, 28.27, 27.88, 27.7, 27.5, 27.06, 25.6, 22.15, 21.87, 13.95; **FT-IR** ($\tilde{\nu}$ cm<sup>-1</sup>): 3858.46, 3740.92, 3609.77, 3322.70, 3052.42, 2950.07, 2784.54, 2628.61, 2405.51, 2303.15, 1968.90, 1828.16, 1720.21, 1529.09, 1452.33, 1381.96, 1238.83, 1174.86, 1022.12, 847.00, 767.04, 713.46, 633.49, 588.72; **HRMS** *m/z* Calcd for C<sub>45</sub>H<sub>65</sub>N<sub>5</sub>O<sub>14</sub>Na 922.4426, Found 922.4423.

#### Synthesis of HBS-constrained single α-helical turn 13<sub>7</sub>3 analogue (11a):

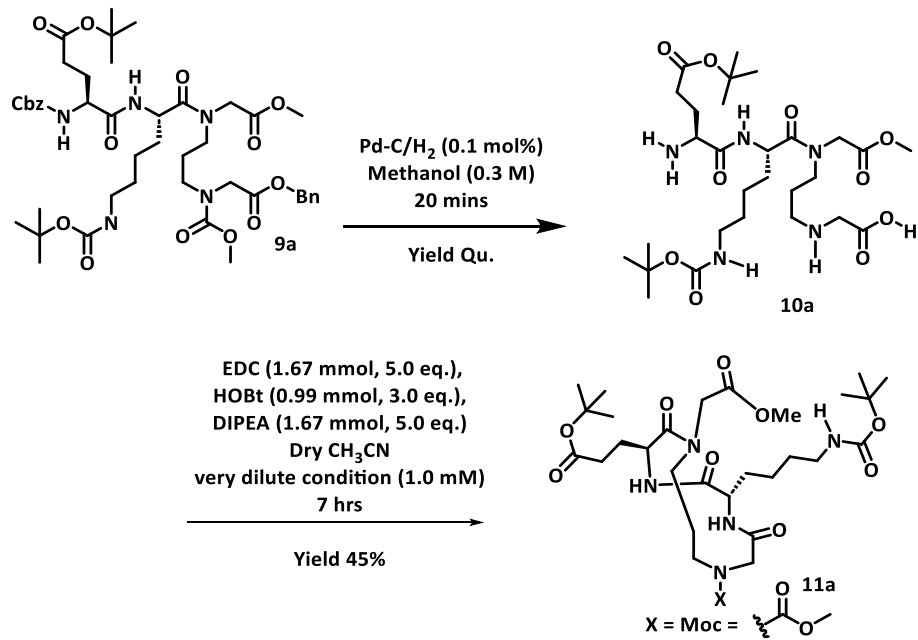

MeOH (3.0 ml) was added to a mixture of **9a** (300 mg, 0.33 mmol) and Pd-C (0.1 mol%) in a sealed round-bottomed flask kept under $\text{H}_2$-atmosphere and stirred for 20 mins by which time TLC indicated the complete consumption of starting material. Filtering the mixture through Whatman-40 filter paper and concentrating the organic filtrate gave the N- and C-terminal double-deprotected derivative of **9a**, **10a**, as a viscous liquid (225.25 mg, 0.33 mmol). Next **10a** was dissolved in dry acetonitrile (1.0 mM) (333.3 ml) followed by addition of EDC (318.3 mg, 1.67 mmol), HOBT (134.9 mg, 0.99 mmol) and DIPEA (282.9 $\mu\text{l}$, 1.67 mmol) under $\text{N}_2$ atmosphere. The mixture was stirred for further 7 hrs. Removal of solvent resulted in a residue which was dissolved in DCM (30.0 ml) and washed with water (2 X 15.0 ml), 1N HCl (2 X 15.0 ml) and saturated $\text{NaHCO}_3$ (2 X 15.0 ml). The organic layer was dried over anhyd. $\text{Na}_2\text{SO}_4$ and concentrated to get a residue which was subjected to purification by silica gel flash column chromatography (EtOAc:Hexane – 9.5:0.5) to yield the desired product as a white solid (98.7 mg, 0.15 mmol, 45% yield); (TLC- EtOAc:Methanol (9:1) – $R_f$ = 0.5); $^1\text{H NMR}$ (400 MHz, $\text{CDCl}_3$, 10 mM) $\delta$ ppm: major conformation: 7.72 (s, 1H, $\text{H}^{\text{N}}\text{Glu}$), 7.26 (s, 1H, $\text{H}^{\text{Nside chain}}\text{Lys}$ merged with $\text{CDCl}_3$), 6.25 (s, 1H, $\text{H}^{\text{N}}\text{Lys}$), 4.67–4.55 (m, 2H, $\text{H}^{\alpha}\text{Lys}$ + $\text{H}^{\alpha}\text{Gly}_1$), 4.49 – 4.39 (m, 2H, $\text{H}^{\alpha}\text{Gly}_1$ + $\text{H}^{\alpha}\text{Gly}_4$), 4.28 – 4.17 (m, 1H, $\text{H}^{\alpha}\text{Gly}_4$), 4.12 – 4.09 (m, 1H, $\text{H}^{\alpha}\text{Glu}$), 3.76 (s, 3H, $\text{H}^{\text{Me}}\text{OMe}$), 3.72 (s, 3H, $\text{H}^{\text{Me}}\text{Moc}$), 3.66 – 3.55 (m, 2H, $\text{H}^{\omega}\text{Lys}$), 3.41 – 3.34 (m, 1H, $\text{Hprp}^c$), 3.09 (bs, 1H, $\text{Hprp}^a$),

2.62 – 2.48 (m, 2H, H<sup>γ</sup>Glu + H<sup>α</sup>Prp<sup>c</sup>), 2.40 -2.33 (m, 1H, H<sup>γ</sup>Glu ), 2.01 (bs, 2H, H<sup>β</sup>Glu), 1.84 – 1.76 (m, 4H, H<sup>α</sup>Prp<sup>b</sup> + H<sup>β</sup>Lys + H<sup>δ</sup>Lys), 1.62 – 1.59 (m, 1H, H<sup>δ</sup>Lys), 1.32 – 1.25 (m, 3H, H<sup>γ</sup>Lys + H<sup>α</sup>Prp<sup>b</sup>) ; <sup>13</sup>C NMR (100 MHz, CDCl<sub>3</sub>, 60 mM) δ ppm: (mixtures of major and minor conformers) 173.7, 173.3, 172.97, 172.0, 170.3, 169.2, 157.9, 156.5, 110.5, 81.8, 65.9, 57.2, 53.0, 52.7, 52.5, 52.2, 48.5, 48.2, 46.5, 43.4, 40.6, 34.5, 32.5, 30.9, 29.7, 29.5, 28.5, 28.0, 25.3, 24.1, 22.7, 15.3 ; **FT-IR** ($\tilde{\nu}$ cm<sup>-1</sup>): 3564.19, 3309.90, 2944.69, 2844.51, 2407.10, 2291.95, 1689.82, 1542.69, 1453.13, 1369.96, 1249.22, 1162.06, 1008.52, 825.41, 728.65, 534.34; **HRMS** *m/z* Calcd for C<sub>30</sub>H<sub>51</sub>N<sub>5</sub>O<sub>11</sub>Na 680.3483, Found 680.3480.

#### Synthesis of C-terminal extended Cyclic peptides (ERG HM-2):

The synthesis of the cyclic peptidomimetic ERG-HM was performed using standard Fmoc solid-phase peptide synthesis (SPPS) on Rink Amide resin in a peptide synthesis syringe. Fmoc deprotection was achieved by treating the resin with 20% piperidine in DMF for 5 min, draining, and washing with DMF (2–3 ×), followed by a second incubation with 20% piperidine in DMF for 15 min. The resin was thoroughly washed with DMF (5 ×) until all traces of residual piperidine were removed. Each canonical amino acid coupling was carried out using Fmoc-protected amino acid (2.0 equiv.), activated *in situ* with HOBt (2.5 equiv.) and N, N'-diisopropylcarbodiimide (DIPC, 2.5 equiv.) in anhydrous DMF for 3 h.

To enable site-specific fluorophore conjugation, Lys15 was incorporated as Fmoc-Lys(Alloc)-OH. Following the assembly of the linear peptide chain, the final *N*-terminal coupling was executed using the free carboxylic acid derivative of the macrocyclic building block, Moc-[GE(OTBu)K(Boc)]G-OH (2.0 equiv.), activated identically with HOBt (2.5 equiv.) and DIPC (2.5 equiv.) in DMF for 3 h.

Prior to global resin cleavage, the Alloc protecting group on the ε-amino group of Lys15 was selectively removed under mild conditions. The resin was washed with anhydrous DCM and treated with a degassed solution of CHCl<sub>3</sub>/AcOH/NMM (37:2:1, v/v/v) containing tetrakis(triphenylphosphine)palladium(0) (Pd(PPh<sub>3</sub>)<sub>4</sub>, 3.0 equiv.) for 8hrs. The resin was subsequently washed sequentially with CHCl<sub>3</sub> (3 ×), DMF (3 ×), and DCM (3 ×). The liberated ε-amino group of Lys15 was then directly labeled by coupling with Rhodamine B (2.0 equiv.), pre-activated with HOBt (2.5 equiv.) and DIPC (2.5 equiv.) in DMF for 3 h at room temperature.

Following successful fluorophore incorporation, the resin was extensively washed with DMF and DCM and dried *in vacuo*. Global cleavage and simultaneous side-chain deprotection were performed by treating the resin with a cleavage cocktail of TFA/H<sub>2</sub>O/TIPS/(95:2.5:2.5) for 3.5 h at room temperature. The cleavage mixture was concentrated under a stream of nitrogen, and the crude peptide was precipitated by adding ice-cold diethyl ether, isolated by centrifugation, and dried. The crude material was purified via reverse-phase HPLC, and the molecular identity and purity were confirmed by MALDI-TOF MS: calculated for [M+H] **5032.867**, found **4990.682** [M-42]<sup>+</sup> (loss of carbodiimide from Arg) and **5070.691** [M+K-H]<sup>+</sup>.

**Maldi Tof (ERG HM-2):**

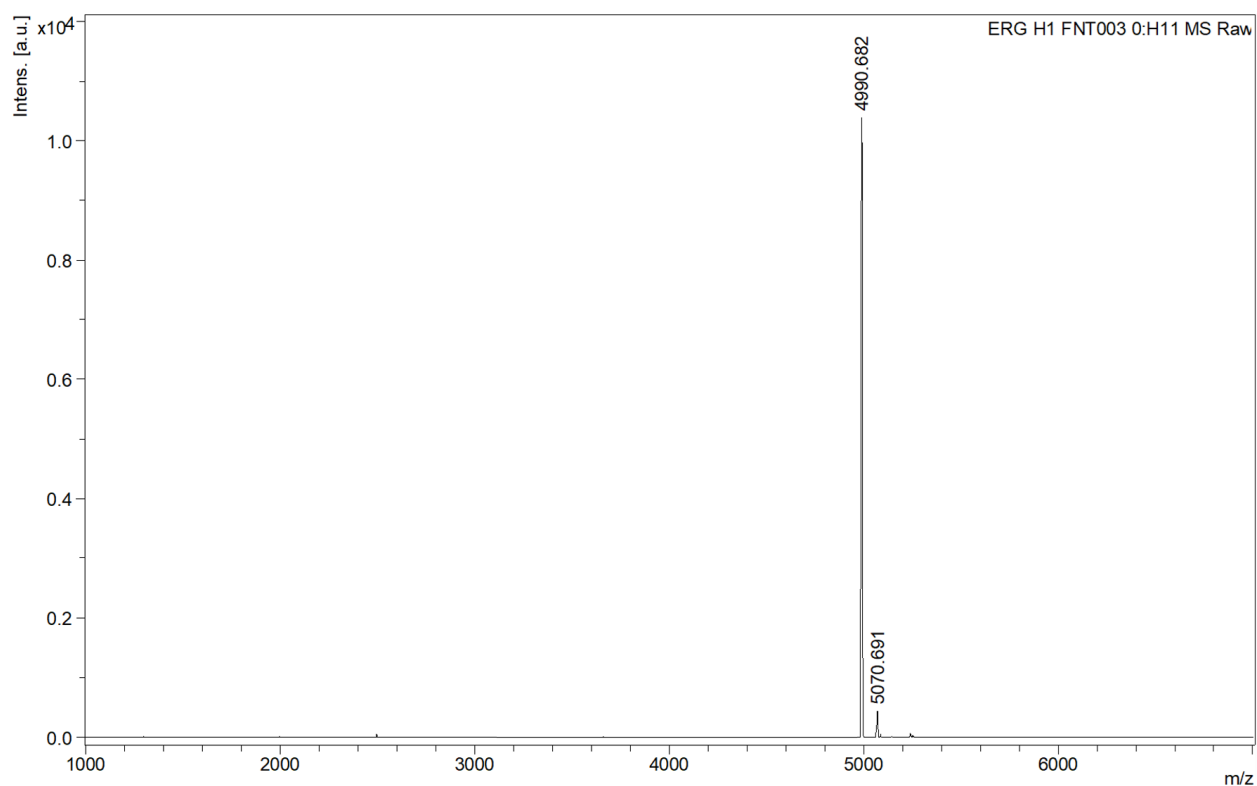

### HPLC chromatogram:

#### <Chromatogram>

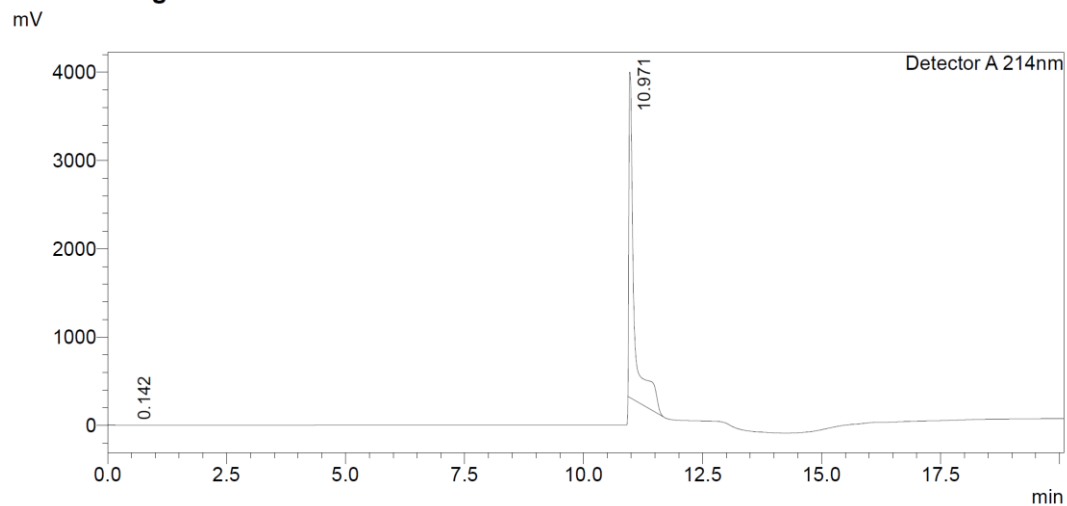

**Figure S4. Analytical RP-HPLC chromatograms and HRMS characterization of ERG HM-2.** Representative analytical RP-HPLC traces (C18 column, $\lambda = 214$ nm, flow rate = 1.0 mL/min; gradient: 0–5 min, 5% CH<sub>3</sub>CN; 5–35 min, linear gradient up to 80% CH<sub>3</sub>CN; 35–39 min, 100% CH<sub>3</sub> CN in H<sub>2</sub>O; mobile phases: solvent A = H<sub>2</sub>O with 0.06% TFA, solvent B = CH<sub>3</sub>CN with 0.05% TFA). Corresponding high-resolution mass spectra (HRMS) confirm the purity and target monoisotopic masses of all synthesized peptidomimetic variants.

#### Solid-Phase Peptide Synthesis of Acyclic Control Peptides (Control peptide and ERG HM-1):

Control peptide and ERG HM-1 were assembled via manual SPPS on Rink Amide AM resin (0.10 mmol, loading: 0.661 mmol/g). Iterative Fmoc deprotections used 20% piperidine/DMF (2 × 10 min). Amino acid couplings employed standard Fmoc-protected residues (2.0 equiv.), HOBt (2.5 equiv.), and DIPC (2.5 equiv.) in DMF for 1.0–1.5 h. Fluorophore incorporation at Fmoc-Lys(Alloc)-OH proceeded via selective side-chain deprotection using Pd(PPh<sub>3</sub>)<sub>4</sub> (3.0 equiv.) dissolved in CHCl<sub>3</sub>/AcOH/NMM (37:2:1) for 8 h, followed by coupling with Rhodamine B (2.0 equiv.) with HOBt (2.5 equiv.) and DIPC (2.5 equiv.) in DMF for 12 h. After final *N*-terminal Fmoc removal, capping was performed with Ac<sub>2</sub>O (2.5 equiv.) and DIPC (2.5 equiv.) in DMF (2 × 15 min). Concomitant resin cleavage and global deprotection were executed using TFA/H<sub>2</sub>O/TIPS/(95:2.5:2.5) for 3.5 h. Crude peptides were precipitated in cold Et<sub>2</sub>O, centrifuged (4000 rpm, 10 min), and purified by semi-preparative RP-HPLC (C18 column, MeCN/H<sub>2</sub>O containing 0.05% and 0.06% TFA, respectively, flow rate: 3.0 mL/min over 45 min).

Lyophilization afforded the control peptides (>95% purity), with identities confirmed by analytical RP-HPLC and MALDI-TOF MS (**Fig. S4**).

**Control peptide:** ToF MS; calculated for $[M + H]^+$ $m/z$ 4366.4498 found $m/z$ 4366.968 $[M+H]^+$.

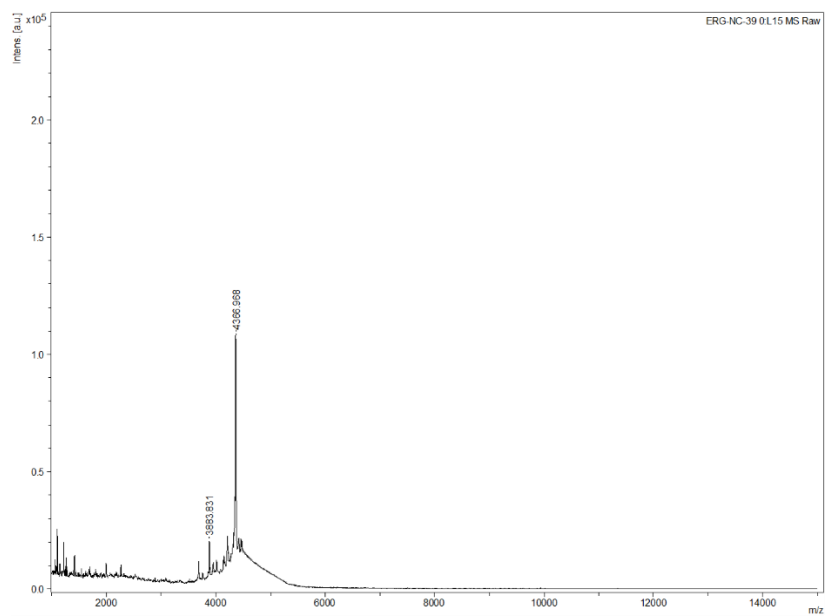

#### HPLC Chromatogram:

<Chromatogram>

mV

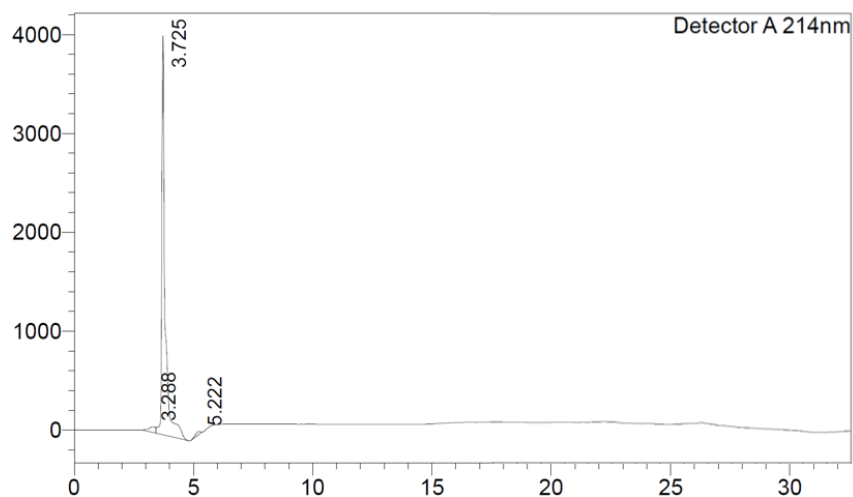

**Figure S5. Analytical RP-HPLC chromatograms and HRMS characterization of Control peptide.** Representative analytical RP-HPLC traces (C18 column, $\lambda = 214$ nm, flow rate = 1.0 mL/min; gradient: 0–5 min, 5% $\text{CH}_3\text{CN}$; 5–35 min, linear gradient up to 80% $\text{CH}_3\text{CN}$; 35–39

min, 100% CH<sub>3</sub>CN in H<sub>2</sub>O; mobile phases: solvent A = H<sub>2</sub>O with 0.06% TFA, solvent B = CH<sub>3</sub>CN with 0.05% TFA). Corresponding high-resolution mass spectra (HRMS) confirm the purity and target monoisotopic masses of all synthesized peptidomimetic variants.

##### ERG HM-1:

#### LC-MS:

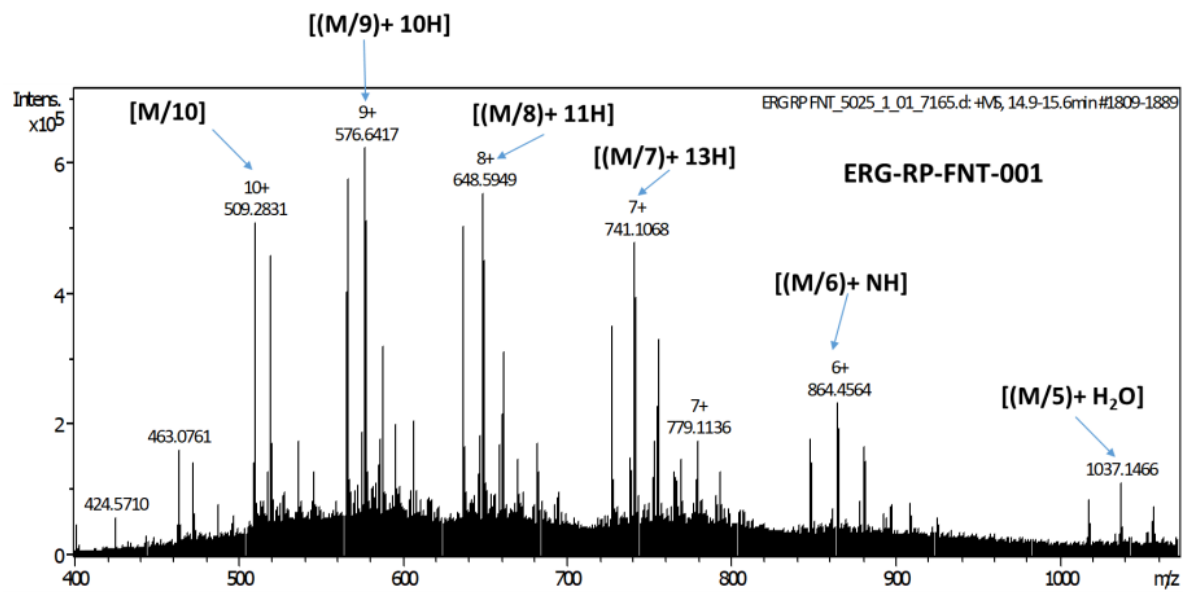

##### HPLC Chromatogram:

**Figure S6. Analytical RP-HPLC chromatograms and HRMS characterization of ERG HM-1.** Representative analytical RP-HPLC traces (C18 column, $\lambda = 214$ nm, flow rate = 1.0 mL/min; gradient: 0–5 min, 5% CH<sub>3</sub>CN; 5–35 min, linear gradient up to 80% CH<sub>3</sub>CN; 35–39 min, 100% CH<sub>3</sub>CN in H<sub>2</sub>O; mobile phases: solvent A = H<sub>2</sub>O with 0.06% TFA, solvent B = CH<sub>3</sub>CN with

**<sup>1</sup>H NMR of Boc-Glu(O-tBu)-Lys(Boc)-OMe (b) in CDCl<sub>3</sub> (10 mM, 400 MHz):**

**$^{13}\text{C}$ NMR of Boc-Glu(O-tBu)-Lys(Boc)-OMe (b) in $\text{CDCl}_3$ (60 mM, 100 MHz):**

**$^1\text{H}$ NMR (400 MHz) of 2a in $\text{CDCl}_3$ (10 mM):**

$^{13}\text{C}$ NMR (100 MHz) of 2a in $\text{CDCl}_3$ (60 mM):

$^1\text{H}$ NMR (400 MHz) of 4a in $\text{CDCl}_3$ (10 mM):

$^{13}\text{C}$ NMR (100 MHz) of 4a in $\text{CDCl}_3$ (60 mM):

$^1\text{H}$ NMR (400 MHz) of 7a in $\text{CDCl}_3$ (10 mM):

**$^{13}\text{C}$ NMR (100 MHz) of 7a in $\text{CDCl}_3$ (60 mM):**

**$^1\text{H}$ NMR (400 MHz) of 9a in $\text{CDCl}_3$ (10 mM):**

**$^{13}\text{C}$ NMR (100 MHz) of 9a in $\text{CDCl}_3$ (60 mM):**

**$^1\text{H}$ NMR (400 MHz) of 11a in $\text{CDCl}_3$ (10 mM):**

**$^{13}\text{C}$ NMR (100 MHz) of 11a in $\text{CDCl}_3$ (60 mM):**

### References:

1. Luscombe NM, Laskowski RA, Thornton JM. Amino acid-base interactions: a three-dimensional analysis of protein-DNA interactions at an atomic level. *Nucleic Acids Res* **2001**;29:2860-74
2. Tam JZ, Palumbo T, Miwa JM, Chen BY. Analysis of Protein-Protein Interactions for Intermolecular Bond Prediction. *Molecules* **2022**;27
